# Structural and functional network-level reorganization in the coding of auditory motion directions and sound source locations in the absence of vision

**DOI:** 10.1101/2021.07.28.454106

**Authors:** Ceren Battal, Ane Gurtubay-Antolin, Mohamed Rezk, Stefania Mattioni, Giorgia Bertonati, Valeria Occelli, Roberto Bottini, Stefano Targher, Chiara Maffei, Jorge Jovicich, Olivier Collignon

## Abstract

hMT+/V5 is a region in the middle occipito-temporal cortex that responds preferentially to visual motion in sighted people. In case of early visual deprivation, hMT+/V5 enhances its response to moving sounds. Whether hMT+/V5 contains information about motion directions and whether the functional enhancement observed in the blind is motion specific, or also involves sound source location, remains unsolved. Moreover, the impact of this crossmodal reorganization of hMT+/V5 on the regions typically supporting auditory motion processing, like the human Planum Temporale (hPT), remains equivocal. We used a combined functional and diffusion MRI approach and individual in-ear recordings to study the impact of early blindness on the brain networks supporting spatial hearing, in male and female humans. Whole-brain univariate analysis revealed that the anterior portion of hMT+/V5 responded to moving sounds in sighted and blind people, while the posterior portion was selective to moving sounds only in blind participants. Multivariate decoding analysis revealed that the presence of motion directions and sound positions information was higher in hMT+/V5 and lower in hPT in the blind group. While both groups showed axis-of-motion organization in hMT+/V5 and hPT, this organization was reduced in the hPT of blind people. Diffusion MRI revealed that the strength of hMT+/V5 – hPT connectivity did not differ between groups, whereas the microstructure of the connections was altered by blindness. Our results suggest that the axis-of-motion organization of hMT+/V5 does not depend on visual experience, but that blindness alters the response properties of occipito-temporal networks supporting spatial hearing in the sighted.

**Significance Statement:** Spatial hearing helps living organisms navigate their environment. This is certainly even more true in people born blind. How does blindness affect the brain network supporting auditory motion and sound source location? Our results show that the presence of motion directions and sound positions information was higher in hMT+/V5 and lower in hPT in blind relative to sighted people; and that this functional reorganization is accompanied by microstructural (but not macrostructural) alterations in their connections. These findings suggest that blindness alters crossmodal responses between connected areas that share the same computational goals.

## INTRODUCTION

In everyday life, moving events are often perceived simultaneously across vision and audition. The brain must therefore exchange and integrate visual and auditory signals to create a unified representation of motion. In the visual system, hMT+/V5, a region in the middle occipito-temporal cortex has long been known for its preferential tuning to visual motion (Watson *et al*., 1993; Tootell *et al*., 1995) and its axis-of-motion columnar organization that supports visual direction selectivity in human and non-human primates (Albright et al., 1984; Diogo et al., 2003; Zimmermann et al., 2011). In the auditory system, the planum temporale (hereafter hPT), engages preferentially in the processing of moving sounds (Baumgart *et al*., 1999; Krumbholz *et al*., 2005). Similar to what was found in hMT+/V5 for the processing of visual directions, hPT also codes for the direction of sounds following an axis-of-motion organization (Battal *et al*., 2019). In addition to their similar coding organization, hMT+/V5 and hPT are part of a network involved in the processing and integration of audio-visual motion information (Gurtubay-Antolin *et al*., 2021). For instance, in addition to its well-documented role for visual motion, hMT+/V5 also responds preferentially to auditory motion (Poirier *et al*., 2005) and contains information about planes of motion using a similar representational format in vision and audition (Rezk *et al*., 2020). Examining how audio-visual motion networks develop in people with no functional vision presents a unique opportunity to assess how changes in sensory experience shape multisensory networks.

In case of early visual deprivation, hMT+/V5 shows enhanced response to auditory motion (Poirier *et al*., 2006; Wolbers, et al., 2011; Jiang, Stecker and Fine, 2014; Dormal *et al*., 2016). Which part of the hMT+/V5 show selectivity to auditory motion in both sighted and blind individuals and which part reorganizes in the early blind remains however debated. Studies in sighted people have shown that the anterior portion of hMT+/V5, called MTa, shows motion selectivity in both vision and audition (Battal et al., 2019, Rezk et al., 2020). It is therefore possible that blindness triggers a posterior extension of this multisensory motion selective region, but this was never formally tested. Moreover, whether hMT+/V5 selectively involves in the processing of auditory motion or whether this region engages for spatial hearing in general remains unknown.

The impact of early acquired blindness on temporal regions typically coding for sounds remains controversial. Some studies suggested an expansion of tonotopic areas and increased response to sounds in the temporal cortex of blind people (Elbert *et al*., 2002; Gougoux *et al*., 2009) while others suggested a reduced involvement of hPT region for auditory motion (Dormal *et al*., 2016; Jiang *et al*., 2016).

Recent advances in multivariate pattern analyses provide the opportunity to go beyond the observation of motion selectivity in a brain region but also probe for the presence of directional information encoded in a region, and whether the region codes information following an axis of motion organization (Albright et al., 1984, Rezk et al., 2020, Battal et al., 2019). No study so far however investigated whether the axis of motion organization known to be implemented in hPT and hMT+/V5 is altered in early blind people.

Previous studies in non-human primates (Palmer and Rosa, 2006; Majka *et al*., 2019) and recent work in humans (Gurtubay-Antolin *et al*., 2021) provided evidence for the putative existence of direct occipito-temporal structural connections between visual and auditory motion selective regions, which may support the exchange of spatial/motion information across audition and vision. The impact of visual deprivation on these connections remains unexplored.

The present study aimed to functionally and structurally investigate how early blindness affects brain networks involved in spatial hearing. Our first aim was to clarify whether and how auditory motion direction and sound source location are represented in the hMT+/V5 and hPT in both sighted and blind individuals. Second, relying on diffusion-weighted imaging, we investigated whether early blindness affects the macro- and microstructure of hMT+/V5 – hPT white matter connections.

## MATERIALS AND METHODS

### Participants

Sixteen early blind (EB) and 18 sighted control (SC) participants were recruited for the study. Participants were matched for age and gender. Sighted participants also participated in an independent visual motion localizer task. One SC participant was excluded due to poor performance on the task within the scanner and another SC participant was excluded due to excessive motion during the scanning session. This resulted in a total of 32 participants included in the analyses: 16 early blind participants (8 female, age range: 20 to 46, mean ± standard deviation (SD) = 33.7 ± 7.2 years) and 16 sighted participants (8 female, age range: 20 to 42, mean ± SD = 31.8 ± 5.7 years). An additional 17 sighted participants (10 females, age range: 20 to 41, mean ± SD = 28 ± 5.3 years) participated in an independent auditory motion localizer experiment. Thirteen of the 16 early blind participants (7 female, mean ± SD = 31.2 ± 5.4 years) and 12 of the 16 sighted participants (7 female, mean ± SD = 32.4 ± 6.2 years) underwent diffusion-weighted magnetic resonance imaging (dMRI). Additionally, 3 sighted participants that did not conduct any functional localizer task were included in the diffusion analyses.

In all cases, blindness was attributed to peripheral deficits with no additional neurological problems (for characteristics of blind participants, see Table 1). All the blind participants lost sight since birth or had visual problems since birth that evolved toward complete blindness before 4 years of age. Seven blind participants had faint light perception without pattern or color vision. Sighted participants had normal or corrected-to-normal vision. Experiments were undertaken with the understanding and written consent of each subject. All the procedures were approved by the research ethics boards of the Centre for Mind/Brain Sciences (CIMeC) and the University of Trento, and in accordance with The Code of Ethics of the World Medical Association, Declaration of Helsinki (Rickham, 1964).

**Table 1.**
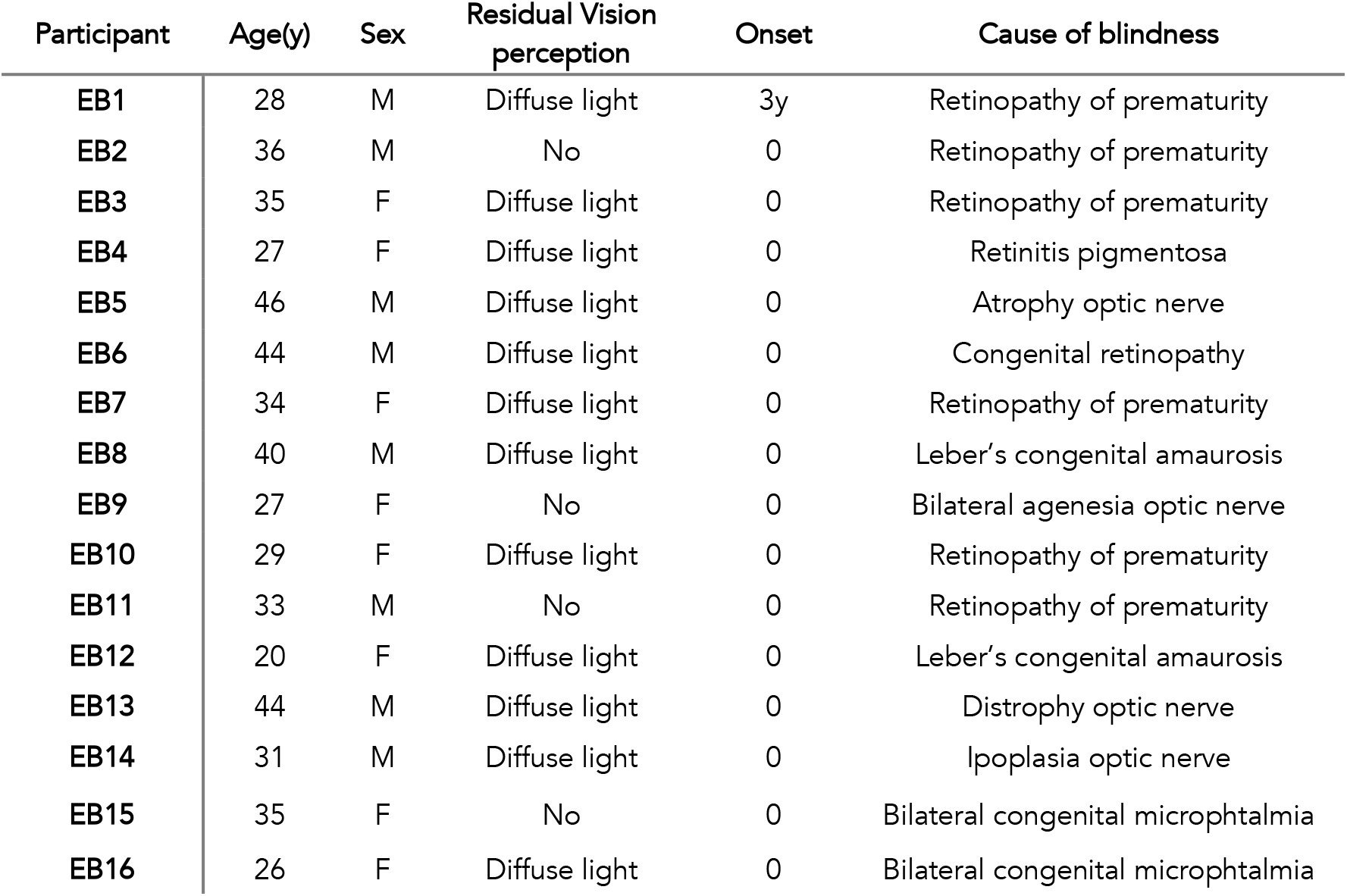
Characteristics of the blind participants. Handedness was evaluated using an adapted version of the Edinburgh inventory, Blind and sighted participants were classified as musicians if they had practiced a musical instrument or had vocal training for at least 2 years on a regular basis (at least 2 hours a week), A: Ambidextrous, M: male, F: female, y: years

The functional and structural data from sighted participants have been published in two previous studies (Battal et al., 2019; Gurtubay et al., 2020). The present study addresses a separate question: how the brain networks supporting spatial hearing change in case of congenital blindness by acquiring new data on this population. The sighted data used in previous work here mostly serve as a control group to be compared with blind people.

### Experimental Design

#### Auditory stimuli

To infer the direction or location of sounds in the horizontal plane of the head, the auditory system extracts interaural differences in arrival time and sound level (ITDs and ILDs, respectively; Middlebrooks and Green, 1991; Blauert, 1997). Up-down localization notably relies on the interaction of sound waves within the pinnae, resulting in idiosyncratic direction-dependent spectral acoustic filters (Musicant and Butler, 1984; Wightman and Kistler, 1989; Middlebrooks and Green, 1992; Hofman, Van Riswick and Van Opstal, 1998). To create an externalized ecological sensation of sound source location and motion (Møller *et al*., 1996), we relied on individual in-ear stereo recordings that were recorded in a semi-anechoic room and from 30 loudspeakers on horizontal and vertical planes, mounted on two semicircular wooden structures with a radius of 1.1m (see Fig 1A) (Battal *et al*., 2020). Binaural in-ear recordings allow binaural properties such as interaural time and intensity differences, as well as participant-specific monaural filtering cues, and serve to create reliable and ecological auditory space (Pavani *et al*., 2002). Participants were seated in the center of the apparatus with their head on a chinrest, such that the speakers on the horizontal plane were at the participant’s ear level and those on the vertical plane were aligned with the participant’s mid-sagittal plane.

**Figure 1.**
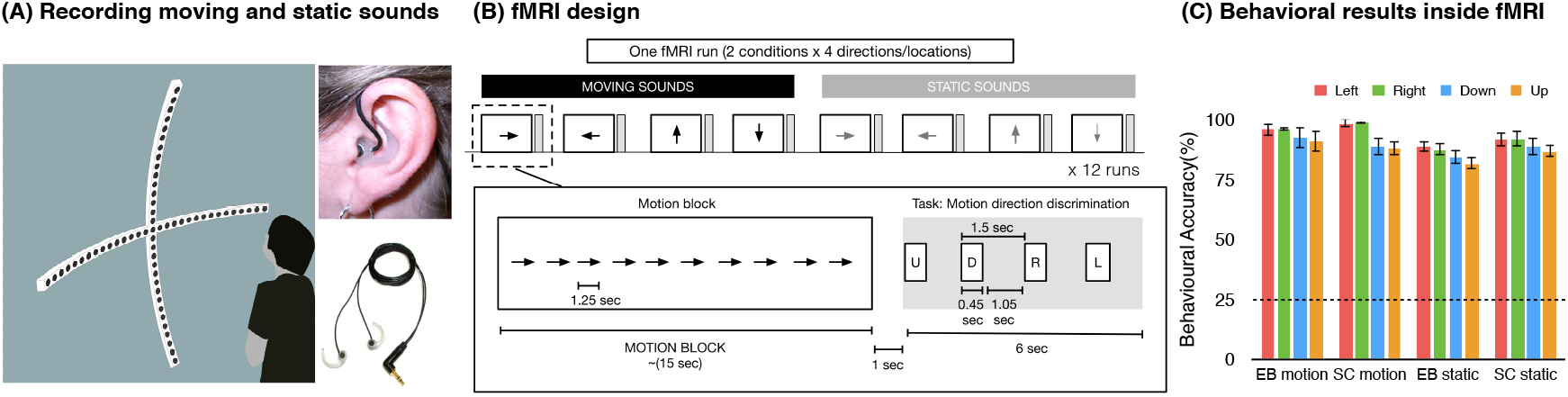
Stimuli and Experimental Design. **(A).** Acoustic apparatus used to present auditory moving and static sounds while binaural recordings were carried out from each participant’s ear before the fMRI session. **(B).** Auditory stimuli presented inside the fMRI consisted of 8 conditions: leftward, rightward, downward, and upward moving sounds and left, right, down, and up static sounds. **(C).** The behavioral performance was recorded inside the scanner; dashed line indicates the chance level.

Auditory stimuli were prepared using a custom-designed MATLAB script (R2013b, MathWorks). During the presentation of stimuli, the audio was recorded using binaural in-ear omni-directional microphones (Sound Professionals-TFB-2; ‘flat’ frequency range 20–20,000 Hz) attached to a portable Zoom H4n digital wave recorder (16-bit, stereo, 44.1 kHz sampling rate). Microphones were positioned at the opening of participant’s left and right auditory ear canals. Then, these recordings were re-played to the participants when they were inside the functional MRI (fMRI). By using in-ear recordings, auditory stimuli automatically convolved with each individuals’ own pinna and head related transfer function to produce a salient auditory perception in external space. The recorded auditory stimuli were used in both the main auditory experiment and the auditory motion localizer. All participants wore a blindfold throughout the auditory recordings and were instructed to keep their eyes closed. Prior to the recordings, the sound pressure level (SPL) was measured from the subject’s head position and ensured that each speaker conveys 65dB-A SPL.

#### Stimuli recordings

Sound stimuli consisted of 1250 ms pink noise (50 ms rise/fall time). In the motion condition, the same pink noise was presented moving in 4 directions: leftward, rightward, upward, and downward. Moving stimuli covered 120° of space/visual field in horizontal and vertical axes. To create the perception of smooth motion, the 1250 ms of pink noise was fragmented into 15 equal length pieces with each 83.333 ms fragment being played every two speakers, and moved one speaker at a time, from the outer left to the outer right (rightward motion), or vice-versa for the leftward motion. For example, for the rightward sweep, sound was played through speakers located at −60° and −52° consecutively, followed by −44°, and so on. A similar design was used for the vertical axis. This resulted in participants perceiving moving sweeps covering an arc of 120° in 1250 ms (speed, 96°/s; fade in/out, 50 ms) containing the same sounds for all four directions. The choice of the movement speed of the motion stimuli aimed to create a listening experience relevant to conditions of everyday life. Moreover, at such velocity it has been demonstrated that human listeners are not able to make the differences between concatenated static stimuli from motion stimuli elicited by a single moving object (Poirier et al., 2017), supporting the participant’s report that our stimuli were perceived as smoothly moving (no perception of successive snapshots). In the static condition, the same pink noise was presented separately at one of the following four locations: left, right, up, and down. Static sounds were presented at the second-most outer speakers (−56° and +56° in the horizontal axis, and +56° and -56° in the vertical axis) to avoid possible reverberation differences at the outermost speakers. The static sounds were fixed at one location at a time instead of presented in multiple locations (Krumbholz et al., 2005; Smith et al., 2004, 2010; Poirier et al., 2017). This strategy was purposely adopted for two main reasons. First, randomly presented static sounds can evoke a robust sensation of auditory apparent motion (Strybel and Neale, 1994; Lakatos and Shepard, 1997; for review, see Carlile and Leung, 2016). Second, presenting static sounds located in a given position and moving sounds directed toward the same position allowed us to investigate whether moving and static sounds share a common representational format using multidimensional scaling (see below), which would have been impossible if the static sounds were randomly moving in space.

Participants were instructed to listen to the stimuli, without performing any task. Stimuli recordings were conducted in a session that lasted approximately 10 minutes, requiring the participant to remain still during this period. All participants reported strong sensation of auditory motion and were able to detect directions and locations with high accuracy during the scanner session (see Fig 1C).

### Auditory experiment

Participants completed a total of 12 runs. Auditory stimuli were presented via MR-compatible closed-ear headphones (Serene Sound, Resonance Technology), and the amplitude was adjusted according to each participant’s comfort level. To familiarize the participants with the task, they completed a practice outside of the scanner while lying down until they reached above 80% of accuracy. All participants wore a blindfold with sterile gauze on top of the eyelids during the auditory task and were instructed to keep the eyes closed during the entire duration of the experiment. Each run consisted of the 8 conditions (4 motion and 4 static), randomly presented using a block-design. Each condition was presented for 15 s block (12 repetitions of each event of 1250 ms sound, no ISI) and followed by 7 s gap in which the participant had to indicate the corresponding direction/location in space and 8s of silence (total inter-block interval was 15 s). The ramp applied at the beginning and at the end of each sound creates static bursts and prevented adaptation to the static sounds. During the response gap, participants heard a voice saying “left”, “right”, “up”, and “down” in pseudo-randomized order. Participants were asked to press a button with their right index finger when the auditory block’s direction or location was matching with the auditory cue (Fig 1B). The number of targets and the order (position 1-4) of the correct button press were balanced across conditions. This procedure was adopted to ensure that the participants gave their response using equal motor command for each condition and to ensure the response is produced after the end of the stimulation period. Each scan consisted of one block of each condition, resulting in a total of 8 blocks per run, with each run lasting 4 m 10 s (100 volumes). The order of the blocks was pseudo-randomized within each run, and across participants.

### Auditory localizer

To localize regions responding to auditory motion, an independent group of sighted participants (n = 17) undertook an auditory motion localizer scan. Individual in-ear recordings of moving and static stimuli were presented in a blocked design. Each block contained 12 repetitions of 1200 ms sounds from one of 8 conditions: 4 motion directions, and 4 static locations. Stimuli within a block were separated by 100 ms ISIs, and each block was followed by a 6 s rest period. The localizer had one run and consisted of 13 repetitions of each condition block in a pseudorandom order. The scan lasted a total of 9 min and 48 s (235 volumes). Participants were instructed to indicate via button press with their right index finger when they detected a stimulus with a shorter duration (targets = 600 ms). The number of targets in each block was varied between 1 and 3 targets, with the location in the block randomized and balanced across conditions. Participants were familiarized with the task before the fMRI session and were blindfolded throughout the scan.

### Visual hMT+/V5 localizer

To identify hMT+/V5, sighted participants undertook an independent visual motion localizer scan. Visual stimuli were back-projected onto a screen (width: 42 cm, frame rate: 60 Hz, screen resolution: 1024 x 768 pixels; mean luminance: 109 cd/m^2^ via a liquid crystal projector (OC EMP 7900, Epson Nagano) positioned at the back of the scanner and viewed via mirror mounted on the head coil at a distance of 134 cm. Stimuli were 16 s of random-dot patterns, consisting of circular aperture (radius 4°) of radial moving and static dots (moving and static conditions, respectively) with a central fixation cross (Huk, Dougherty and Heeger, 2002). One hundred and twenty white dots (diameter of each dot was 0.1 visual degree) were displayed on a grey background, moving 4° per second. In all conditions, each dot had a limited lifetime of 0.2 s. Limited lifetime dots were used in order to ensure that the global direction of motion was determined by integrating local signals over a larger summation field rather than by following a single dot (Bex, Simmers and Dakin, 2003). Stimuli were presented for 16 s followed by a 6 s rest period. Stimuli within motion blocks alternated between inward and outward motion (expanding and contracting) once per second. Because the localizer aimed to localize the global hMT+/V5 complex (e.g., including both MT and MST subregions) the static block was composed of dots maintaining their position throughout the block to prevent flicker-like motion (Smith 2006). The localizer consisted of 14 alternating blocks of moving and static dots (7 each) and lasting a total of 6 m 40 s (160 volumes). To maintain the participant’s attention and to minimize eye-movement during acquisition throughout the localizer’s run, participants were instructed to detect a color change (from black to red) of a central fixation cross (0.03°) by pressing the response button with the right index finger.

### Imaging parameters

Functional and structural data were acquired with 4T Bruker MedSpec Biospin MR scanner, equipped with 8-channel head coil. Functional images were acquired with T2*-weighted gradient echo-planar sequence with fat suppression. Acquisition parameters included a repetition time of 2500 ms, echo time of 26 ms, flip angle of 73°, a field of view of 192 mm, a matrix size of 64 x 64, and voxel size of 3 x 3 x 3 mm. A total of 39 slices were acquired in ascending feet-to-head interleaved order with no gap. The three initial scans of each acquisition run were discarded to allow for steady-state magnetization. Before each EPI run, we performed an additional scan to measure the point-spread function (PSF) of the acquired sequence, including fat saturation, which served for distortion correction that is expected with high-field imaging (Zeng and Constable, 2002).

High-resolution anatomical scan (Papinutto and Jovicich, 2008) was acquired using a T1-weighted 3D MP-RAGE sequence (176 sagittal slices, voxel size of 1 × 1 × 1mm; field of view 256 x 224 mm; repetition time = 2700 ms; TE = 4.18 ms; FA: 7°; inversion time: 1020 ms). Participants were blindfolded and instructed to lie still during acquisition. Foam padding was used to minimize scanner noise and head movement. Diffusion weighted images were acquired using an EPI sequence (TR = 7100 ms, TE = 99 ms, image matrix = 112 × 112, FOV = 100 × 100 mm^2^, voxel size 2.29 mm isotropic). Ten volumes without any diffusion weighting (b0-images) and 60 diffusion-weighted volumes with a b-value of 1500 s/mm^2^ were acquired.

### Univariate fMRI analysis

Raw functional images were pre-processed and analyzed with SPM8 (Welcome Trust Centre for Neuroimaging London, UK; http://www.fil.ion.ucl.ac.uk/spm/software/spm/) implemented in MATLAB R2014b (MathWorks). Before the statistical analysis, our preprocessing steps included slice time correction with reference to the middle temporal slice, realignment of functional time series, coregistration of functional and anatomical data, spatial normalization to an echo planar imaging template conforming to the Montreal Neurological Institute space, and spatial smoothing (Gaussian kernel, 6 mm FWHM).

#### Auditory experiment

To obtain blood oxygen level-dependent (BOLD) activity related to auditory spatial processing, we computed single subject statistical comparisons with fixed-effect general linear model (GLM). In the GLM, we used eight regressors from each condition (four motion directions, four sound source locations). The canonical double-gamma hemodynamic response function implemented in SPM8 was convolved with a box-car function to model the above mentioned regressors. Motion parameters derived from realignment of the functional volumes (3 translational motion and 3 rotational motion parameters), button press, and four auditory response cue events were modeled as regressors of no interest. During the model estimation, the data were high-pass filtered with cut-off 128 s to remove the scanner drift and low-frequency fluctuations from the time series. To account for serial correlation due to noise in fMRI signal, autoregressive (AR (1)) was used.

At the fixed-effect individual subject level (FFX), to obtain activity related to auditory processing in the whole brain, the contrasts tested the main effect of each condition: Left Motion, Right Motion, Up Motion, Down Motion, Left Static, Right Static, Up Static, and Down Static. Next, to identify regions responding preferentially to the auditory motion and static stimuli, we compared the response of all motion conditions to all static conditions (Motion > Static, and Static > Motion). These linear contrasts generated statistical parametric maps (SPM[T]) that were further spatially smoothed (Gaussian kernel 8 mm FWHM) before being entered in a second-level group analysis, using a random effect model (RFX), accounting for inter-subject variance.

At the group level, a series of one-sample t-tests was implemented to examine the main effects of each condition (Motion, Static) and motion processing (Motion > Static) for each group. A conjunction analysis isolated brain areas jointly activated for the contrast Motion > Static in both groups (EB and SC). Two-sample t-tests were then performed to compare these effects between groups (SC > EB, EB > SC).

Statistical inferences were done using family-wise error (FWE) correction for multiple comparisons using *p<0.05* over the entire brain volume or over small spherical volumes (15 mm radius) located around regions of interest (see Table 1) using a minimal cluster size threshold of 20 contiguous voxels (Worsley *et al*., 1996). Significant clusters were anatomically labeled using the xjView Matlab toolbox (http://www.alivelearn.net/xjview) or structural neuroanatomy information provided in the Anatomy Toolbox (Eickhoff *et al*., 2007).

### Independent visual and auditory motion localizer

#### Region of interest definition

We used independent auditory and visual motion localizers to functionally defined hPT and hMT+/V5 regions. Preprocessing steps were similar to whole-brain univariate analysis (see section *Univariate fMRI Analysis*). Single subject statistical comparisons were made using a fixed-effect GLM for each participant with two regressors (motion, static), and motion parameters (6 regressors of no interest). The canonical double-gamma hemodynamic response function implemented in SPM8 was convolved with a box-car function for each regressor. Motion parameters derived from realignment of the functional volumes (3 translational motion and 3 rotational motion parameters); button press was modeled as regressor of no interest. During the model estimation, the data were high-pass filtered with cut-off 128 s to remove the scanner drift and low-frequency fluctuations from the time series. To account for serial correlation due to noise in fMRI signal, autoregressive (AR (1)) was used.

One-sample t-tests were conducted to characterize the main effect of motion processing (Motion > Static). This linear contrast generated statistical parametric maps that were further spatially smoothed (Gaussian kernel 8 mm FWHM) and entered a second-level group analysis using a random effects GLM. Group-level peak coordinates of bilateral hMT+/V5 and hPT were defined by contrasting the main effects of localizer scan (Motion vs Static), surviving a whole-brain family-wise-error correction (p<0.05). Peak coordinates from the auditory and visual motion localizers were used to create a sphere of 6 mm radius (110 voxels) around 4 regions-of-interest (ROIs): left hMT+/V5, right hMT+/V5, left hPT, and right hPT. The 4 ROIs were defined functionally but constrained by anatomical landmarks of the regions. hPT was selected within the triangular region lying caudal to the Helschl’s gyrus on the supratemporal plane, whilst hMT+/V5 was constrained with the ascending limb of the inferior temporal sulcus (Zeki *et al*., 1991; Watson *et al*., 1993).

To prevent the possible spurious overlap between the visual and auditory responsive regions arising from smoothing (Jiang et al., 2015), we performed the statistical inferences on the beta parameter estimates that were extracted from the unsmoothed data.

Because defining the visual motion area hMT+/V5 is obviously impossible in EB, we opted for the peak coordinates from the visual motion localizers in sighted subject in normalized in MNI to be applied to blind and people similarly. We, then, extracted data from the same spatially aligned ROIs in both groups to perform functional univariate and multivariate analyses.

### Multivariate pattern analyses

To investigate the presence of auditory motion direction and sound source location information, multivariate pattern analyses (MVPA) were conducted within the independently defined hMT+/V5 and hPT regions (see “localizers” sections above). All further analyses were conducted on these regions for all sighted and blind participants.

#### Multi-class Direction and Location Decoding

To investigate motion direction and static location information in areas hMT+/V5 and hPT in sighted and blind participants, 4-classes classifiers were trained and tested to discriminate between the response patterns of the 4 auditory motion directions and 4 sound source locations, respectively.

Preprocessing steps were identical to the steps performed for univariate analyses, except for functional volumes that were smoothed with a Gaussian kernel of 2 mm (FWHM). MVPA were performed in CoSMoMVPA (http://www.cosmomvpa.org/; Oosterhof et al. 2016, which implements LIBSVM software (http://www.csie.ntu.edu.tw/~cjlin/libsvm)). A general linear model was implemented in SPM8, where each block was defined as a regressor of interest. A t-map was calculated for each block separately. Two multi-class linear support vector machine (SVM) classifiers with a linear kernel with a fixed regularization parameter of C = 1 were trained and tested for each participant separately within each group. The two multi-class classifiers were trained and tested to discriminate between the response patterns of the 4 auditory motion directions and locations, respectively.

For each participant, the classifier was trained using a cross-validation leave-one run-out procedure where training was performed with n-1 runs and testing was then applied to the remaining run. In each cross-validation fold, an ANOVA-based feature selection was applied on the training data to select a subset of voxels (n = 110) that showed the most significant variation between the categories of stimuli (in our study, between orientations). The selected features were used for training and testing. This feature selection process not only ensures a similar number of voxels within a given region across participants, but, more importantly, identifies and selects voxels that are carrying the most relevant information across categories of stimuli (Cox and Savoy, 2003; De Martino *et al*., 2008), therefore minimizing the chance to include voxels that carry noises unrelated to our categories of stimuli. The t-maps in the training set were normalized (z-scored) across conditions, and the estimated parameters were applied to the test set. To evaluate the performance of the classifier and its generalization across all the data, the previous step was repeated 12 times where in each fold a different run was used as the testing data and the classifier was trained on the other 11 runs. For each region per subject, a single classification accuracy was obtained by averaging the accuracies of all cross-validation folds.

#### Binary Direction and Location Decoding

To investigate the preference of “axis of motion/space” in both hMT+/V5 and hPT, binary classifiers were used to discriminate brain activity patterns for motion direction within and across axes. Four binary classifiers were used to discriminate brain activity patterns for left vs right motion, left vs right static, up vs down motion, up vs down static (hereafter called within-axis classification). We used eight additional classifiers to discriminate across axes (left vs up, left vs down, right vs up, and right vs down motion directions; left vs up, left vs down, right vs up, and right vs down sound source locations; hereafter called across-axes classification).

To assess whether hMT+/V5 and hPT regions demonstrate axis of motion characteristic tuning for auditory motion, we averaged the 2 within-axes motion accuracies from leftwards vs. rightward, and upward vs. downward binary classifications. We also averaged the motion accuracies from 4 across-axes motion binary classification. To assess the existence of axis-of-space, we performed averaging on the sound source location classification accuracies with 2 within-axes and 4 across-axes binary classifications to obtain 1 accuracy value per subject per axis.

#### Multi-dimensional Scaling

To visualize the pairwise decoding accuracies of motion directions and sound source locations, the 28 decoding accuracy values were represented in the form of dissimilarity matrix. Each column and each row of the matrix represented one motion direction or sound location. The matrix elements were the pairwise decoding accuracies. The accuracy values were used as dissimilarity measure. We used multidimensional scaling (MDS) to project the high-dimensional dissimilarity matrix space onto two dimensions with the pairwise decoding that were obtained from hMT+/V5 and hPT in both sighted and blind individuals. The 32 neural dissimilarity matrices (1 per participant) for each of the four ROIs were used as neural input for MDS visualization. Note that MDS are for visualization purpose only and were not used for statistical inferences.

#### Statistical Analysis

Statistical analyses were performed in MATLAB (for non-parametric tests) and R (for repeated-measures ANOVAs, and LMMs). Tests for normality were carried out using a Shapiro-Wilk test. All the within-group comparisons were carried out using paired *t*-tests.

Statistical significance in the multivariate classification analyses was assessed using non-parametric tests permuting condition labels and bootstrapping (Stelzer, Chen and Turner, 2013). Each permutation step included shuffling of condition labels and re-running the classification, which was repeated 100 times on the single-subject level. Next, we applied a bootstrapping procedure to obtain a group-level null distribution that is representative of each group. For each group, from each subject’s null distribution one value was randomly chosen and averaged across all the subjects. This step was repeated 100,000 times resulting in a group level null distribution of 100,000 values. The classification accuracies across subjects were considered as significant if the p<0.05 after corrections for multiple comparisons using the FDR method (Benjamini and Yekutieli, 2001). The group comparison was also tested for significance by using permutation (100,000 iterations).

Classification accuracies were entered into a linear mixed-effects model (LMM), as computed through the *lmer* function in R (afex package, Singmann et al., 2021). We performed a LMM with Group (EB, SC), Condition (motion, static), Region (hMT+/V5, hPT), and Hemisphere (left, right) as fixed effects and subject as a random effect.

To assess axis of motion preference across groups and across ROIs, the pairwise decoding accuracies were submitted to LMMs. For each ROI, we performed a LMM with Group (EB, SC), Condition (within-axis, across-axis), and Hemisphere (left, right) as fixed effects and subject as a random effect.

### Diffusion data analysis

#### Preprocessing of diffusion data

Data preprocessing was implemented in MRtrix 3.0 (Tournier, Calamante and Connelly, 2012) (www.mrtrix.org), and in FSL 5.0.9 (https://fsl.fmrib.ox.ac.uk/fsl/fslwiki/FSL). Data were denoised (Veraart et al., 2016), corrected for Gibbs-ringing, Eddy currents distortions and head motion (Andersson and Sotiropoulos, 2016), and for low-frequency B1 field inhomogeneities (Tustison et al., 2010). Spatial resolution was up-sampled by a factor of 2 in all three dimensions (1.15 mm isotropic) using cubic b-spline interpolation, and intensity normalization across subjects was performed by first deriving scale factors from the median intensity in select voxels of white matter, grey matter, and CSF in b = 0 s/mm^2^ images, and then applying these across each subject image (Raffelt, Tournier, Rose, et al., 2012). This step normalizes the median white matter b = 0 intensity (i.e., non-diffusion-weighted image) across participants so that the proportion of one tissue type within a voxel does not influence the diffusion-weighted signal in another. The T1-weighted structural images were non-linearly registered to the diffusion data in ANTs (Avants et al., 2008) using an up-sampled FA map (1×1×1 mm^3^) and segmented in maps for white matter, grey matter, CSF, and sub-cortical nuclei using the FAST algorithm in FSL (Zhang, Brady and Smith, 2001). This information was combined to form a five tissue type image to be used for anatomically constrained tractography in MRtrix3 (Smith et al., 2012). These maps were used to estimate tissue-specific response functions (i.e. the signal expected for a voxel containing a single, coherently-oriented fiber bundle) for grey matter, white matter, and CSF using Multi-Shell Multi-Tissue Constrained Spherical Deconvolution (MSMT) (CSD) (Jeurissen et al., 2014). Fiber orientation distribution functions (fODFs) were then estimated using the obtained response function coefficients averaged across subjects to ensure subsequent differences in fODFs amplitude will only reflect differences in the diffusion-weighted signal. Note that by using MSMT-CSD in our single-shell, data benefitted from the hard non-negativity constraint, which has been observed to lead to more robust outcomes (Jeurissen et al., 2014). Spatial correspondence between participants was achieved by generating a group-specific population template with an iterative registration and averaging approach using FOD images from all the participants (Raffelt et al., 2011). Each subject’s FOD image was registered to the template using FOD-guided non-linear registrations available in MRtrix (Raffelt, Tournier, Crozier, et al., 2012). These registration matrices were used to transform the seed and target regions from native diffusion space to template diffusion space, where tractography was conducted. We chose to conduct tractography in the template diffusion space, as FOD-derived metrics of microstructural diffusivity can only be computed in that space. Subsequently, we extracted the following quantitative measures of microstructural diffusivity for all the fixels (fiber populations within a voxel) in the brain: fiber density (FD), fiber-bundle cross-section (FC) and a combined measure of fiber density and cross-section (FDC) (Raffelt et al., 2017). For further details on these metrics, see “FOD-derived microstructural metrics” section.

#### Preparation of hMT+/V5 and hPT for tractography

As for the fMRI analyses, hMT+/V5 was functionally defined using the visual motion localizers in the 13 SCs included in the analysis of the diffusion data. To functionally define hPT, we used the peak coordinate of the Motion vs Static contrast in the auditory experiment. To avoid that the location of hPT was biased towards SC, we used the conjunction of the SC and EB participants that underwent the fMRI and dMRI acquisitions.

We computed the warping images between the standard MNI space and the native structural space of each participant by conducting a non-linear registration in ANTs (Gurtubay-Antolin *et al*., 2021). Using these registration matrices, we transformed the group peak-coordinates from the standard MNI space to the native structural space of each participant. We then transformed these coordinates from the native structural space to the native diffusion space using the same transformation used to register the T1-weighted structural images to the diffusion data (described in the Diffusion data preprocessing section). Once in native diffusion space, the peak-coordinates were moved to the closest white matter voxel (FA>0.25) (Blank, Anwander and von Kriegstein, 2011; Benetti *et al*., 2018; Gurtubay-Antolin *et al*., 2021) and a sphere of 5 mm radius was centered there. To ensure that tracking was done from white matter voxels only, we masked the sphere with individual white matter masks. Last, ROIs were transformed from native diffusion space to template diffusion space, where tractography was conducted.

#### Tractography: hMT+/V5 – hPT connections

Probabilistic tractography between hMT+/V5 and hPT was conducted for both hemispheres. In addition to the hPT, a region just anterior to hMT+/V5 (called hMTa; see (Rezk et al., 2020)) is also selectively recruited for the processing of moving sounds (Poirier et al., 2005; Saenz et al., 2008; Battal et al., 2019; Rezk et al., 2020). To avoid that hMT+/V5 – hPT connections could include fibers connecting hPT with hMTa (which lies near hMT+/V5 and responds to auditory motion), individually identified hMTa (as localized by auditory motion localizer task) was used as an exclusion mask.

We computed tractography in symmetric mode (i.e., seeding from one ROI and targeting the other, and conversely). We then merged the tractography results pooling together the reconstructed streamlines. We used two tracking algorithms in MRtrix (‘iFOD2’ and ‘Null Distribution2’). The former is a conventional tracking algorithm, whereas the latter reconstructs streamlines by random tracking. The ‘iFOD2’ algorithm (Second-order Integration over FODs) uses a Bayesian approach to account for more than one fiber-orientation within each voxel and takes as input a FOD image. Candidate streamline paths are drawn based on short curved ‘arcs’ of a circle of fixed length (the step-size), tangent to the current direction of tracking at the current points (Tournier, Calamante and Connelly, 2010). The ‘Null Distribution2’ algorithm reconstructs connections based on random orientation samples, identifying voxels where the diffusion data is providing more evidence of connection than that expected from random tracking (Morris, Embleton and Parker, 2008). We used the following parameters for fiber tracking: randomly placed 5000 seeds for each voxel in the ROI, a step length of 0.6 mm, FOD amplitude cutoff of 0.05 and a maximum angle of 45 degrees between successive steps (Tournier, Calamante and Connelly, 2010, 2012). We applied the anatomically-constrained variation of this algorithm, whereby each participant’s five-tissue-type segmented T1 image provided biologically realistic priors for streamline generation, reducing the likelihood of false positives (Smith et al., 2012). The set of reconstructed connections were refined by removing streamlines whose length was 2.5 SD longer than the mean streamline length or whose position was more than 2.5 SD away from the mean position like previous studies (Gurtubay-Antolin et al., 2021; Yeatman et al., 2012; Takemura et al., 2016). To calculate a streamline’s distance from the core of the tract we resampled each streamline to 100 equidistant nodes and treat the spread of coordinates at each node as a multivariate Gaussian. The tract’s core was calculated as the mean of each fibers x, y, z coordinates at each node.

#### Statistical analysis

Statistical significance in the presence of hMT+/V5 – hPT connections was assessed comparing the number of streamlines reconstructed by random tracking (‘Null Distribution2’ algorithm), with those generated by conventional tracking (‘iFOD2’ algorithm) (Morris, Embleton and Parker, 2008; McFadyen, Mattingley and Garrido, 2019; Gurtubay-Antolin *et al*., 2021). We calculated the logarithm of the number of streamlines [log(streamlines)] to increase the likelihood of obtaining a normal distribution (Müller-Axt, Anwander and von Kriegstein, 2017; Tschentscher et al., 2019), which was tested before application of parametric statistics using the Shapiro–Wilk test in RStudio (Allaire, 2015). The log-transformed number of streamlines were compared using two-sided paired t-tests. To control for unreliable connections, we calculated the group mean and SD of the log-transformed number of streamlines for each connection and we discarded participants whose values were more than 2.5 SDs away from the group mean for the respective connection. Connections were only considered reliable when the number of streamlines reconstructed with the ‘iFOD2’ algorithm were higher than the ones obtained with the ‘null distribution’ algorithm. Significance was thresholded at p = 0.05 Bonferroni-corrected for multiple comparisons (p = 0.025, two hemispheres).

#### Whole tract analysis

Differences between SCs and EBs in hMT+/V5 – hPT connections were assessed using macrostructural and microstructural measures. Macrostructural characterization included the trajectory and the connectivity index, whereas we relied on diffusivity measures derived from fiber orientation distribution for the microstructure assessment.

To assess whether hMT+/V5 – hPT connections followed the same trajectory in the SC and EB groups, we computed their spatial overlap by means of The Dice Similarity Coefficient (DSC) (Dice, 1945). The DSC measures the spatial overlap between regions A and B, and it is defined as DSC (A, B) = 2(A∩B)/(A+B) where ∩ is the intersection. We calculated the DSC of hMT+/V5 – hPT connections, using as region A the sum of binarized tract-density images of hMT+/V5 – hPT connections (thresholded at 6 subjects) in standard space and in the SC group. Region B was the analogous image in the EB group.

The connectivity index was determined by the log-transformed number of streamlines from the seed that reached the target divided by log-transformed the product of the generated sample streamlines in each seed/target voxel (5000) and the number of voxels in the respective seed/target mask (Müller-Axt, Anwander and von Kriegstein, 2017; Tschentscher *et al*., 2019; Gurtubay-Antolin *et al*., 2021).

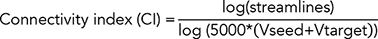

Microstructure assessment relied on the FOD-derived quantitative metrics fiber density (FD), fiber-bundle cross-section (FC) and fiber density and cross-section (FDC) (Fixel-based analysis pipeline in www.mrtrix.org). These estimates characterize microstructure isolating the contribution of crossing fibers within the same voxels of interest, providing more accurate values than tensor-derived metrics such as fractional anisotropy (Behrens *et al*., 2007; Tournier, Calamante and Connelly, 2007; Raffelt *et al*., 2017). Briefly, FD is a quantitative measure of fiber density within a voxel, given that the integral of the FOD along a particular direction is proportional to the intra-axonal volume of axons aligned in that direction. It was calculated as the integral of the FOD along a particular direction and it is sensitive to alterations at the fixel-level (Raffelt et al., 2017). FC reflects changes in a fiber bundle’s intra-axonal volume that are manifested as a difference in the number of spatial extent that the fiber bundle occupies (cross-sectional area). FCs for each fixel were calculated in the plane perpendicular to the fixel direction and were estimated by using the non-linear warps used to register each subject’s FOD to the template. Lastly, multiplying FD and FC we computed the metric FDC, which combines both sources of information (within-voxel FD and changes in FC). These estimates were computed in all the voxels that were crossed by fibers connecting hMT+/V5 and hPT and then averaged for each bundle.

All measures (i.e., connectivity index, FD, FC, FDC) were compared between SCs and EBs using two-sided paired t-tests, after testing for normality using the Shapiro–Wilk test. Significance threshold was set at p = 0.006 (alpha = 0.05 Bonferroni corrected for 2 hemispheres and 4 metrics) and subjects’ values were more than 2.5 SDs away from the group mean were considered outliers. A possible lateralization of group differences in the microstructural metrics was tested by a 2×2 ANOVA with Group as a between-subject factor (2 levels: SC, EB) and Hemisphere as a within-subject factor (2 levels: R, L)

#### Along tract analysis

An along-tract analysis was performed on the reconstructed tracts using the Along Tract Statistics toolbox (Colby et al., 2012) in MATLAB (R2016b). A mean tract geometry (MTG) was estimated for each tract after resampling the streamlines to 50 points, so that all streamlines were composed by the same number of points. For each tract and participant, the mean of diffusion scalar values of corresponding points (i.e., streamline points assigned to the same MTG point) was computed per each MTG point, and individual along-tract profiles were estimated per each tract. To control for the possibility that group differences could be driven by scalar outliers, we excluded tracts whose mean scalar values deviated more than 2.5 SDs from the mean of each subject or deviated for more than 10 consecutive points from the group mean. To further exclude aberrant values, we rejected participants presenting profiles in which at 5 consecutive positions along the tract the profile values deviated ±2.5 SDs for the respective scalar analysis (in a group-wise fashion) (Novello *et al*., 2018).

Per each of the considered diffusion scalars, a linear mixed-effects model was then adopted with fixed effects of group, position (i.e., MTG point), and a group and position interaction to assess possible diffusion scalar changes affecting differently the two groups. A group and position interaction was considered significant for p < 0.008 (alpha = 0.05 Bonferroni-corrected 2 hemispheres and 3 diffusion scalars). We adopted a permutation (n=5000) approach to control the Type 1 error and adjust p-values accordingly: a null distribution of maximum T-values across all MTG points under the null hypothesis of no group differences was empirically estimated by permuting group labels and recording the maximum observed T-statistics. For each MTG point, the group analysis T-statistics was then compared against the null distribution to get an adjusted p-value (Colby et al., 2012). Results were then considered significant for *p* FWE corrected < 0.05.

### Brain-Behavior Correlation Analysis

We investigated the link between behavioral performance and neural activity of hMT+/V5 and hPT regions by performing between-subject Pearson’s correlation on behavioral performance with (1) extracted beta parameter estimates, and (2) extracted decoding accuracies in both EB and SC groups. We also investigated the link between behavioral performance and white matter microstructure by performing between-subject Pearson’s correlation analysis.

Behavioral performance was measured as the accuracy of detecting motion directions and sound source locations during the fMRI session. For every ROI (left and right hemispheres in hMT+/V5 and hPT), we created a sphere of 3-mm radius around the group-level coordinates and extracted the mean beta parameter estimates for auditory motion and static conditions and performed between-subject correlation.

Classification accuracies obtained from multi-class classification from each ROI of each subject was correlated with overall performance of motion direction and sound source location discrimination. Statistical results were corrected for multiple comparisons using the FDR method (Benjamini and Yekutieli, 2001).

## RESULTS

### Behavioral

Behavioral performance in all the 8 conditions in both groups was above 80% of correct responses, demonstrating that we were able to trigger salient and reliable auditory motion/location percepts while the subjects were inside the scanner. To determine if there were any differences between groups or conditions in the target detection task performed during the auditory experiment, accuracy scores were entered into a 2 x 2 x 4 repeated measure ANOVA to test the interaction between Group (EB, SC; between-subject factor), Condition (motion, static; within-subject factor), and Orientation (left, right, up, and down; within-subject factor). Importantly, this showed no main effect of Group (F_1,30_ = 0.401; *p* = 0.5), indicating that the overall accuracy while detecting direction of motion or location of sound source did not differ between the blind and sighted groups. There was a significant main effect of Condition (F_1,30_ = 11.49; *p* = 0.002), which was caused by higher accuracy in the motion condition as compared to the static condition. There was a significant main effect of Orientation (F_1.6,48.3_ = 14.24; *p* < 0.001), caused by greater accuracy in the horizontal orientations (left and right) as compared to the vertical orientations (up and down). Post-hoc two-tailed t-tests (*p* < 0.05, Bonferroni corrected for multiple comparisons) showed that this main effect was due to a significant difference between left orientation with up (t_15_ = 5.22, *p* < 0.001) and down (t_15_ = 3.87, *p* = 0.001) orientations, and between right orientation with up (t_15_ = 5.17, *p* < 0.001) and down (t_15_ = 3.81, *p* = 0.001) orientations. No interaction between Condition x Orientation was observed.

### Univariate analyses

#### Whole brain analyses

To identify brain regions that are preferentially recruited for auditory motion processing in SC and EB groups, we performed a univariate whole brain analysis. Figure 2 shows the response to motion and static auditory stimuli in EB and SC participants. Consistent with previous studies (Pavani *et al*., 2002; Warren, Zielinski and Green, 2002; Poirier *et al*., 2005), a preferential response to auditory moving stimuli (Motion > Static) was observed for SC participants in the superior temporal gyri, bilateral hPT, precentral gyri, and anterior portion of middle temporal gyrus in both hemispheres (Fig 2A). A similar response was observed in EB participants, with a reliable extension toward the occipital cortex (Fig 2B).

**Figure 2.**
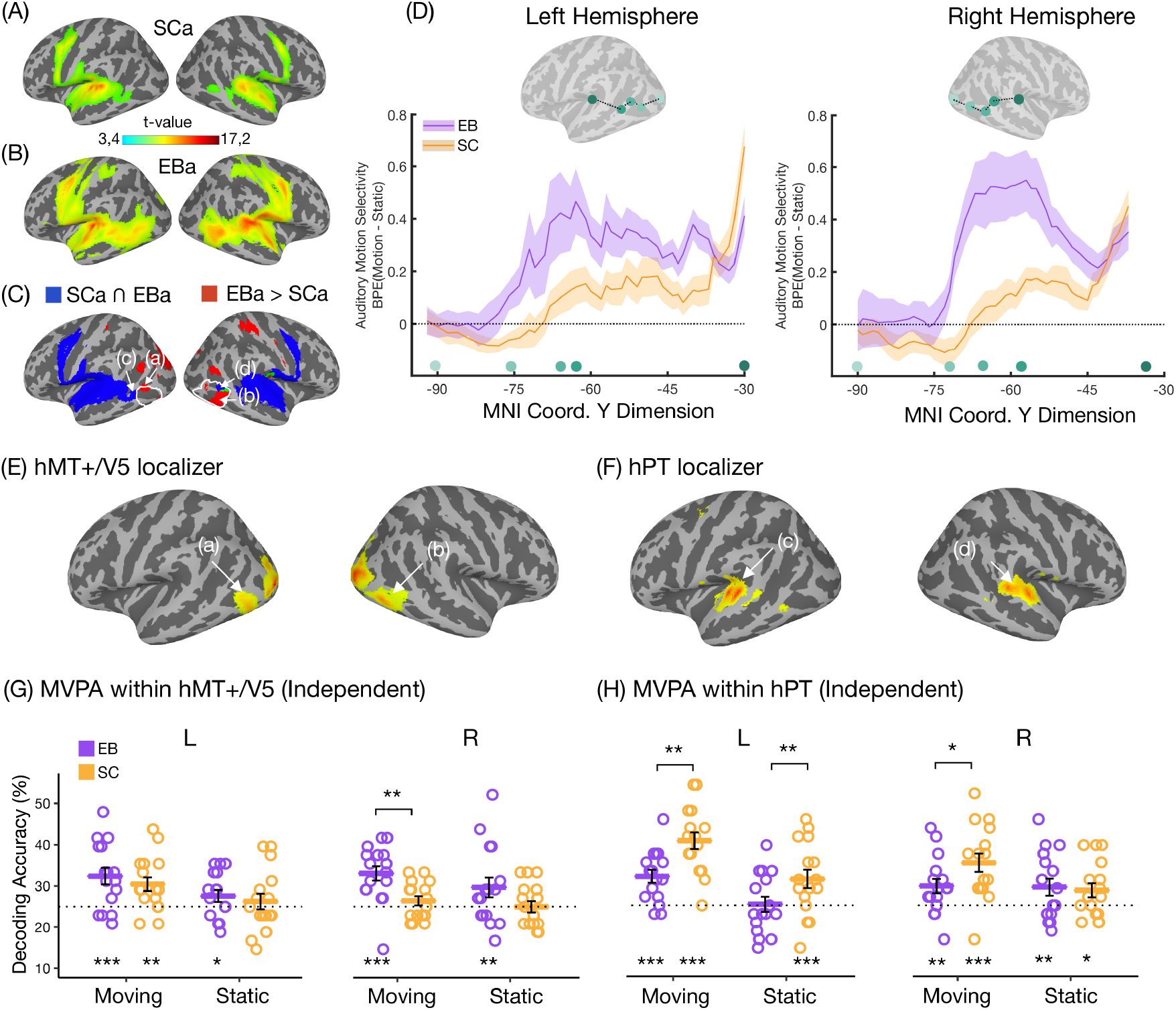
Univariate and multi-class MVPA results. **(A-B).** Activations obtained from the contrast testing which regions preferentially activated for auditory motion processing in sighted (SCa) and early blind (EBa) participants: **(A).** Sighted [Motion > Static], **(B).** Blind [Motion > Static]. **(C).** Activation in blue indicates auditory motion selectivity in both groups [Sighted conj, Blind] x [Motion > Static]. Activation in red indicates enhanced responses to moving compared to static sounds in the early blind compared to the sighted group [Blind > Sighted] x [Motion > Static]. Activation in green indicates the overlap between the conjunction and group comparison analyses. The arrows indicate the peak coordinates of middle temporal gyrus (MTG) in the group comparison ((a) and (b)) and conjunction ((c) and (d)) analyses. All the maps are thresholded with p<0.001 (uncorrected) for illustration purpose only (see methods for statistical significance assessment). **(D).** Auditory motion selectivity (beta parameter estimates (arbitrary units ± SEM) in blind (purple) and in sighted (gray) extracted along the geodesic path between significant peak coordinates from the V1 and hMT+/V5 from visual motion localizer, from auditory main experiment group comparison MTG ((a) and (b)), the group conjunction MTG ((c) and (d)) and the group conjunction hPT. **(E-F).** Motion selective results from the visual and auditory motion localizers in left and right hemispheres (FWE corrected p<0.05). **(G-H).** Decoding accuracies of 4 auditory motion directions and 4 static locations in independently localized in left and right hMT+/V5 and hPT in blind (purple) and in sighted (yellow). Error bars indicate the SEM, the dashed line indicate the chance level (%25). Results are FDR corrected (\**p* < 0.05, \*\**p* < 0.01, \*\*\**p* < 0.001).

To identify regions responding more to moving than static sounds in both EB and SC participants, we ran a conjunction (AND; Nichols et al. 2005) analysis (SC [motion > static] ∩ EB [motion > static]). This showed that both groups activated the superior temporal gyrus, bilateral hPT and the anterior portion of middle temporal gyrus bilaterally. The right middle temporal gyrus (MTG) region partially overlapped with the functionally defined hMT+/V5 identified visually (motion > static) at the group level in SC participants (white outline, Fig 2C).

To identify which regions activated more for moving than static sound in EB versus SC participants, we performed a two-sample t-test (EB [motion > static] > SC [motion > static]). This revealed enhanced activity for EB participants in regions including the precuneus, the cuneus extending into the intraparietal sulci, and bilateral posterior middle temporal gyrus (see Table 2 for whole-brain univariate analyses results).

**Table 2.**
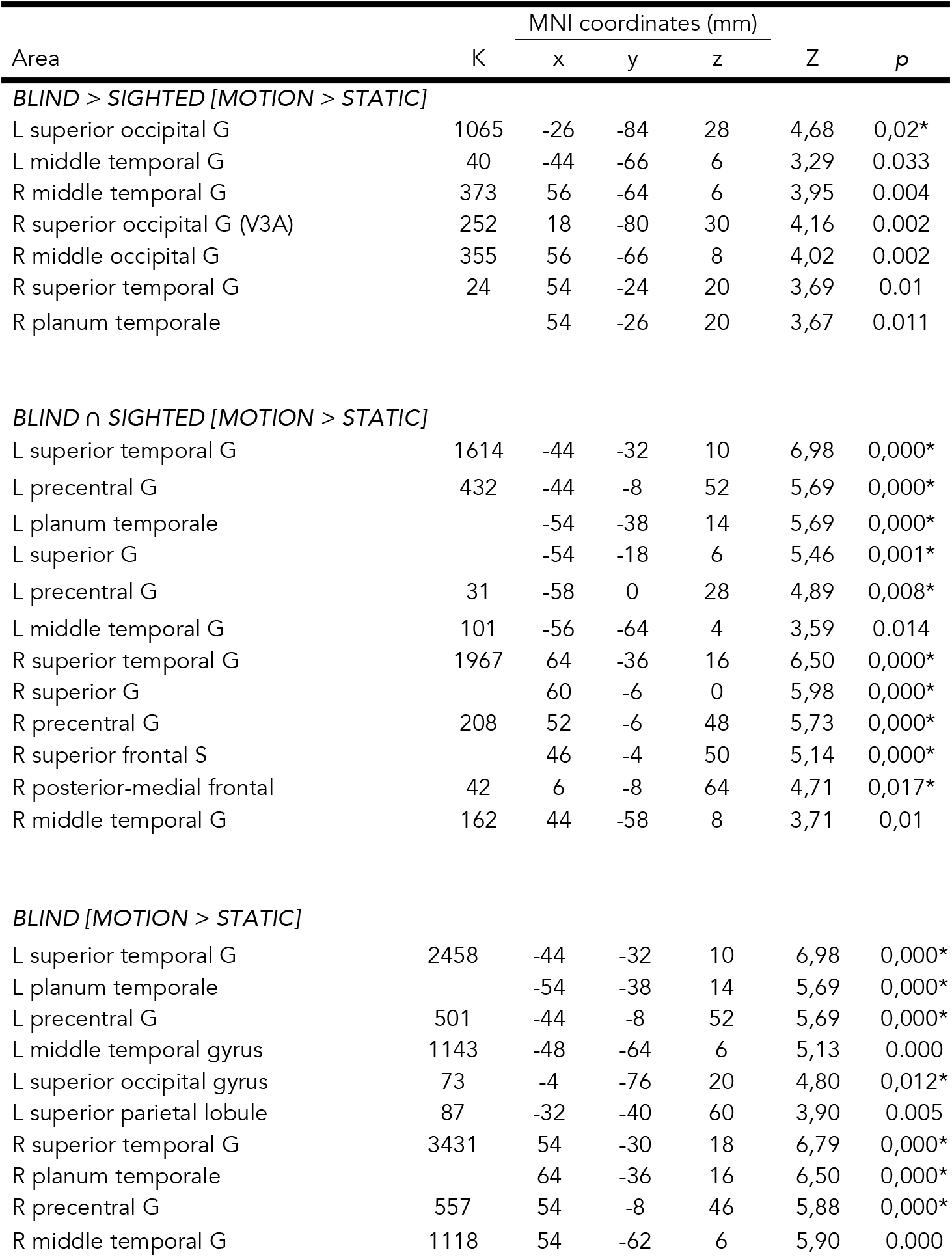
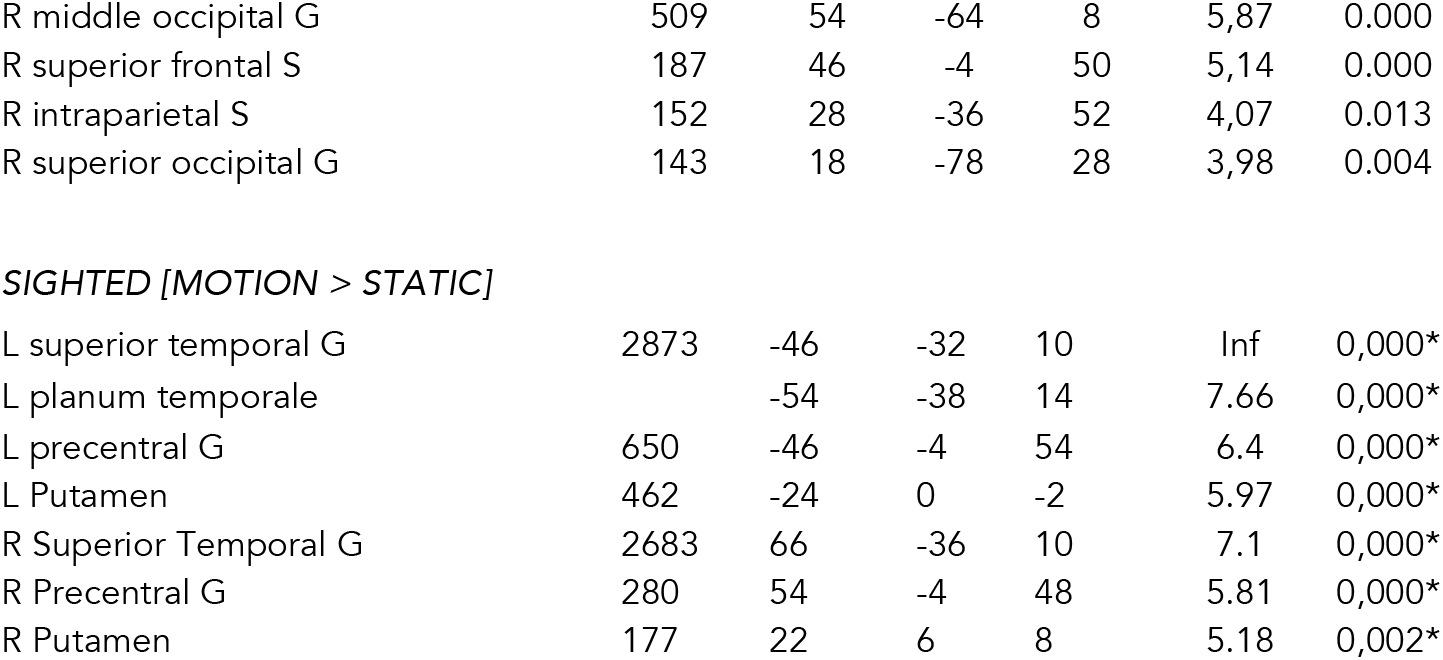
Results of the univariate analyses for the main effect of auditory motion processing [motion > static] in the blind and the sighted. Coordinates reported in this table are significant (p < 0.05 FWE) after correction over small spherical volumes or over the entire brain volume (*). Coordinates used for correction over small spherical volumes were extracted from four papers investigating auditory motion processing in the sighted only (Alink et al., 2012; Pavani et al., 2002) or comparing sighted and blinds (Dormal et al., 2016; Collignon et al., 2009) and are as follows (x, y, z, in MNI space): right hPT [66 −36 14] (Dormal et al., 2016); right middle occipital gyrus [48 −76 6] (Collignon et al., 2011); left middle temporal gyrus (hMT +/V5) [− 42 − 64 4] (Dormal et al., 2016); right middle temporal gyrus (hMT +/V5) [42 − 60 4] (Dormal et al., 2016); K represents the number of voxels when displayed at p(uncorrected) < 0,001, L: left, R: right, G: gyrus, S: sulcus.

#### ROI univariate analysis

##### ROI definition

To avoid circularity that can arise from selection of ROIs, more particularly “double dipping” (Kriegeskorte et al., 2009) – the use of the same dataset for selection and specific analysis – we independently localized visual and auditory motion responsive areas. Whole-brain univariate analyses for independent visual and auditory motion localizers were performed to acquire the peak coordinates of hMT+/V5 and hPT, selective to visual and auditory motion respectively. The obtained stereotactic MNI coordinates were as follows: L hMT+/V5: [-46 −72 −2]; R hMT+/V5: [40 −76 −2], and L hPT: [-48 −30 8], R hPT: [62 −36 10].

To investigate whether auditory motion and static sound elicited responses within ROIs across groups, beta parameters extracted from hPT and hMT+/V5 regions were entered into two separate 2 x 2 x 2 repeated measure ANOVA, Group (EB, SC) as between-subject factor and Hemisphere (left, right), and Condition (motion, static) as within-subject factors.

For the hPT region, we observed a main effect of Condition (F_1,30_ = 148.4, p < 0.001, h^2^ =0.2). The main effect of Condition was caused by a greater response to motion > static stimuli. The interaction of Group x Condition (F_1,30_ = 5.6, p = 0.03, h^2^ =0.01) was significant. Post-hoc two-tailed t-tests (p < 0.05, Bonferroni corrected) showed that the interaction was caused by greater responses to motion over static in SC (t_30_ = 10.29, p < 0.001) than EB group (t_30_ = 6.94, p < 0.001). A significant interaction of Hemisphere x Group (F_1,30_ = 4.7, p = 0.04, h^2^ =0.04) was due to the right hPT in EB group showing higher activity compared to left PT (t_30_ = 1.2, p = 1), while SC group showing higher activity in the left hPT when compared to right hPT (t_30_ = 1.9, p = 0.4; p < 0.05 Bonferroni corrected post-hoc two-tailed t-tests). The interaction between Hemisphere x Condition (F_1,30_ = 20.5, p < 0.001, h^2^ =0.02) was driven by the difference between motion and static conditions was bigger in the left hPT (t_30_ = 11.4, p <0.001) compared to the right hPT (t_30_ =6.77, p <0.001).

For the hMT+/V5 region, we observed an interaction of Group x Condition (F_1,30_ = 4.7, p = 0.04, h^2^ =0.02) showing (p < 0.05, Bonferroni corrected) greater responses to motion over static in the EB (t_30_ = 2.86, p = 0.04). The interaction between Hemisphere x Condition (F_1,30_ = 5.6, p = 0.025, h^2^ =0.03) was driven by increased motion selectivity in the left hemisphere (t_30_ = 2.3, p = 0.057; post-hoc two-tailed t-tests p<0.05 Bonferroni corrected).

As expected from univariate analyses (Battal et al., 2019), beta parameter estimates did not show any evidence for motion direction or sound-source location specific activity.

### ROI multivariate pattern analyses

#### Multi-class Motion Direction and Static Location Decoding

To further investigate the presence of information about auditory motion direction and sound source location, we ran multi-class MVP-decoding in four ROIs identified using independent auditory and visual motion localizers (hMT+/V5 and hPT in both hemispheres). Figure 2F-G shows decoding accuracies for motion (leftward, rightward, upward, downward) and static stimuli (left, right, up, down) in the four regions of interest for EB and SC participants. For motion stimuli, permutation testing (FDR-corrected) revealed that classification accuracies in hMT+/V5 were significantly above chance for EB participants in both hemispheres (left: mean ± SD = 32.4 ± 8, *p* < 0.001; right: mean ± SD = 33.1 ± 6.9, *p* < 0.001). In SC participants, decoding accuracy was significantly above chance in the left hMT+/V5 but not in the right hMT/V5 (left: mean ± SD = 30.5 ± 6.6, *p* = 0.002; right: mean ± SD = 26.4 ± 4.5, *p* = 0.184). In hPT, decoding accuracy was significantly above chance in both hemispheres in both groups (EB left: mean ± SD = 32 ± 6.2, *p* < 0.001; EB right: mean ± SD = 29.7 ± 7, *p* = 0.003; SC left: mean ± SD = 40.6 ± 8, *p* < 0.001; SC right: mean ± SD = 35.3 ± 8.9, p < 0.001). Permutation of two-sample t-tests revealed that decoding accuracy was higher for EB as compared to SC in the right hMT+/V5 (p = 0.02) but not in the left hMT+/V5 (*p* = 0.62). In contrast, decoding accuracy was greater in SC than in EB in the left hPT (*p* = 0.016) but not in the right hPT (*p* = 0.101) (Fig 2G).

For static location stimuli, decoding accuracies were significant within hMT+/V5 in the right (mean ± SD = 29.7 ± 9.6, *p* = 0.003) and very close to the cut-off significance value in the left hemisphere (mean ± SD = 27.6 ± 5.8, *p* = 0.054) of EB participants, while decoding was not significantly greater than chance in either the left or right hMT+/V5 for SC participants (left hMT+/V5: mean ± SD = 26.3 ± 7.6, *p* = 0.2; right hMT+/V5: mean ± SD = 25 ± 5.4, *p* = 0.458). In the hPT, classification accuracy was significantly above chance in both hemispheres in SC participants (left hPT: mean ± SD = 31.3 ± 8.8, *p* < 0.001; right hPT: mean ± SD = 28.7 ± 6.8, *p* = 0.023), but only in the right hemisphere of EB (right hPT: mean ± SD = 29.4 ± 8.3, *p* = 0.007; left hPT: mean ± SD = 25.3 ± 7.3, *p* = 0.458). The decoding accuracy from the two groups were compared using a two-sample t-test, and the statistical significance was assessed using permutations. The results revealed that decoding accuracy for sound source locations was higher for SC as compared to EB in the left hPT (p = 0.002) but not in the right PT (*p* = 0.05).

Finally, we assessed differences between decoding accuracies across groups and regions using an LMM. Group, region, and hemisphere predictors were entered as fixed effects and subjects was entered as a random effect. This revealed a main effect of Condition (F_1,210_ = 26.5, *p* < 0.001) due to higher accuracies for motion over static stimuli across all regions. We also observed a main effect of Region (F_1,210_ = 9, *p* = 0.003) showing overall higher decoding in hPT than hMT+/V5. Crucially, we observed a significant interaction between Group x Region (F_1,30_ = 22.9, *p* < 0.001). Post-hoc two-tailed t-tests (*p* < 0.05; Bonferroni corrected for multiple comparisons) showed that decoding accuracy in hMT+/V5 was significantly greater for EB over SC (t_78.3_ = 2.6, *p* = 0.02), while decoding accuracy in hPT was significantly greater for SC over EB (t_78.3_ = 3.5, *p* = 0.002). The significant interaction between Hemisphere and Group (F_1,210_ = 6.34, p = 0.013) was due to the higher decoding accuracies in the left hemisphere in SC compared to the right hemisphere (t_210_ = 5.5, p < 0.001), and EB group showing marginally higher decoding accuracy in right hemisphere compared to the SC group (t_78.3_ =2.6, p = 0.05), while SC group showing higher decoding accuracy in the left hemisphere compared to EB group (t_78.3_ = 3.5, p = 0.035). The lack of significant interaction between Group x Region x Condition indicates that differences in the decoding accuracies between groups and regions are similar for motion and sound source processing.

#### Axis-of-motion/location preference

To assess the presence of an axis of motion organization, we averaged the binary decoding accuracies of auditory motion directions for each axe type (within and across axes), for each group, and each ROIs. While higher decoding accuracies between pairs of directions (e.g., leftwards vs. upward) indicate that the representation of directions is more distinct, lower decoding accuracies among the directions (e.g., leftwards vs. rightwards) suggests more similar or overlapping representation (Fig 3A-B). The averaged within and across axes binary decoding accuracies (dissimilarity) entered a linear mixed model with group, axes and hemisphere as fixed effects and subject as a random effect.

**Figure 3.**
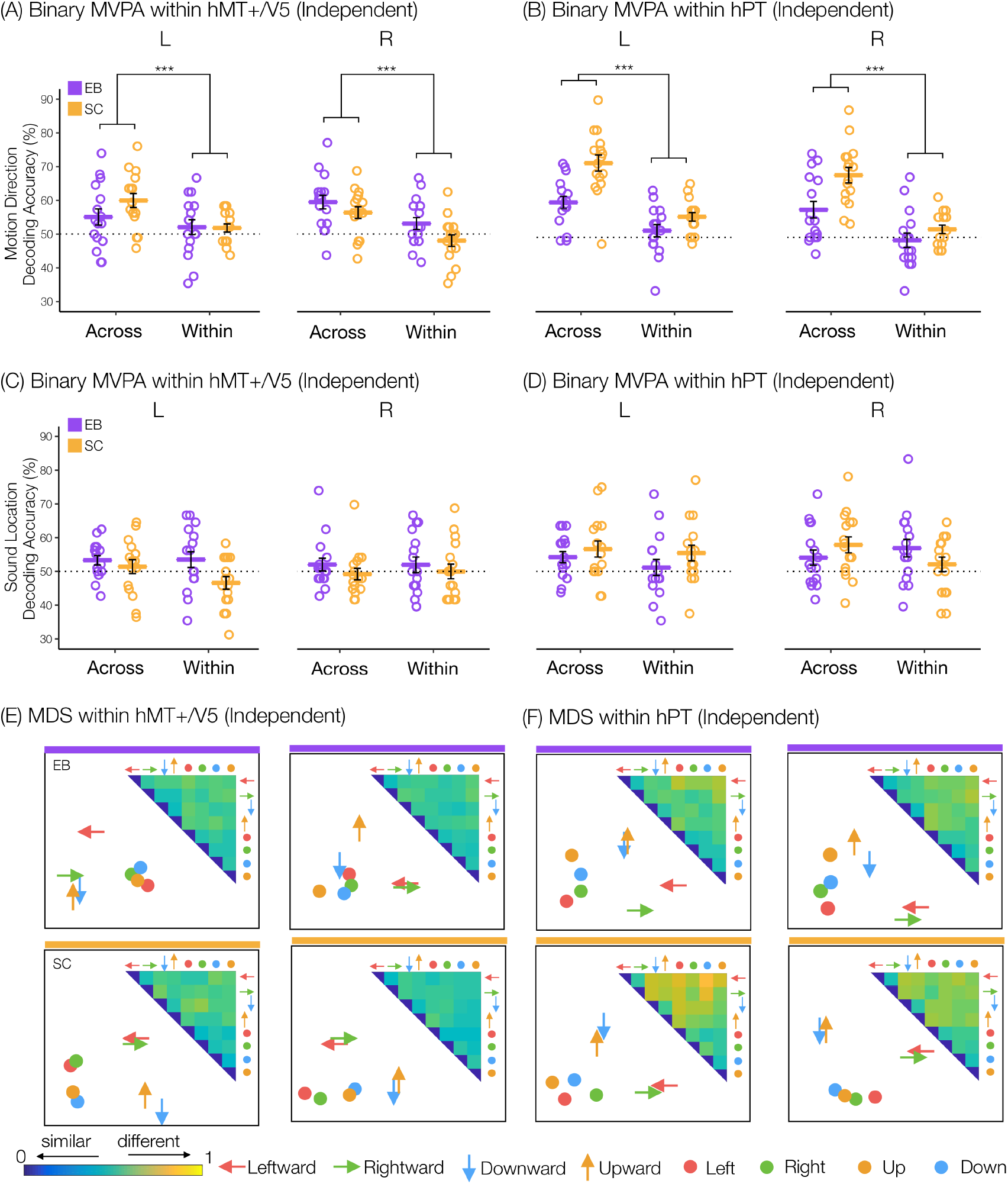
Binary MVPA and multidimensional scaling results. **(A-B).** Averaged across and within axes direction decoding accuracy results in independently localized in left (L) and right (R) hMT+/V5 and hPT in blind (purple) and in sighted (yellow). The asterisks indicate significant main effect of axes that across axes decoding accuracies are higher compared to within axes decoding accuracies (\*\*\**p* < 0.001). **(C-D).** Averaged across and within axes sound source location decoding accuracy results in independently localized in left (L) and right (R) hMT+/V5 and hPT in blind (purple) and in sighted (yellow). Error bars indicate the SEM, the dashed line indicate the chance level (%50). **(E-F).** The inset shows 28 binary decoding accuracy values represented in the form of dissimilarity matrix extracted from left and right hMT+/V5 and hPT in blind (indicated with purple bar) and in sighted (indicated with yellow bar). Binary decoding multi-dimensional scaling (MDS) visualizes the dissimilarity matrices elicited by four motion directions (arrows) and four sound source locations (dots). The pairwise distances between arrows and dots approximately reflect direction/location pattern similarities (dissimilarity measure: accuracy values). While high decoding accuracy values (distinct neural patterns) presented in far apart direction/locations; low decoding accuracy values (similar neural patterns) presented in close together direction/locations. Color codes for arrow/dots are as follows: green indicates left direction/location; red indicates right direction/location; orange indicates up direction/location; and blue indicates down direction/location.

In hPT region, this analysis revealed a significant effect of Axes (F_1,90_ = 84.8, p < 0.001) indicating across-axes directions are more distinct compared to within-axis directions across hemispheres and across groups. We also observed a main effect of Group (F_1,30_ = 22, p <0.001), revealing overall higher binary decoding accuracies in the SC group. The group-by-axes interaction (F_1,90_ = 7.4, p = 0.008) showed that while both group show an axis of motion preference (SC across vs. within axes: t_90_ = 8.4, *p* < 0.0001; EB across vs. within axes: t_90_ = 4.6, *p* = 0.0001), the difference between across and within axes decoding accuracy is higher in sighted than in the blind group (t_76.9_ = 5.3, *p* < 0.0001; *p* < 0.05 Bonferroni corrected for multiple comparisons).

In hMT+/V5 region, we observed a main effect of Axes (F_1,90_ = 25.5, p < 0.001) implying that across-axis directions have higher decoding accuracies compared to within-axis directions across hemispheres and across groups. The significant group-by-hemisphere interaction, (F_1,90_ = 6.3, p = 0.01), was driven by SC group showing higher decoding accuracies in the left hemisphere while EB group showing higher decoding accuracies in the right hemisphere (*p* < 0.05 Bonferroni corrected for multiple comparisons).

LMM on sound source location distances did not reveal significant results involving the groups in both hPT and hMT+/V5 (Fig 3C-D).

#### Multi-dimensional Scaling

We used the binary accuracy values to build neural dissimilarity matrices for each subject and each ROI (Fig 3E-F). Visualization of the binary decoding accuracies of the motion direction and sound source locations further supported a separation between static and moving sounds in both hMT+/V5 and hPT regions and in both groups, and a stronger axis-of-motion preference in SC compared to EB group in the hPT region.

### Diffusion weighted imaging

#### hMT+/V5 – hPT tractography

For the left hemisphere, the number of reconstructed streamlines was significantly above chance in both groups, with the ‘iFOD2’ algorithm generating a significantly higher number of streamlines compared to the ‘Null distribution’ algorithm (SC: log streamlines[iFOD2] = 5.2 ± 1.0, log streamlines[Null distribution] = 3.7 ± 0.8, Paired t-Test, t(14) = 7.7, p = 2 x 10 ^-6^, d=2.0; EB: log streamlines[iFOD2] = 5.2 ± 1.3, log streamlines[Null distribution] = 3.8 ± 1.3, Paired t-Test, t(12) = 3.9, p = 2 x 10 ^-3^, d=1.1). The same results were obtained for right hMT+/V5 – hPT connections (SC: log streamlines[iFOD2] = 4.8 ± 1.1, log streamlines [Null distribution] = 1.4 ± 1.2, Paired t-Test, t(14) = 8.1, p = 1 x 10 ^-6^, d=2.1; EB: log streamlines[iFOD2] = 4.2 ± 1.1, log streamlines[Null distribution] = 1.6 ± 0.9, Paired t-Test, t(12) = 6.8, p = 2 x 10 ^-5^, d=1.9). hMT+/V5 – hPT connections in a representative EB participant as well as group-averaged structural pathways between hMT+/V5 and hPT for the EB group can be seen in Figures 4A and 4B (the single-subject data for the EB group available at https://osf.io/7w8gu/).

**Figure 4.**
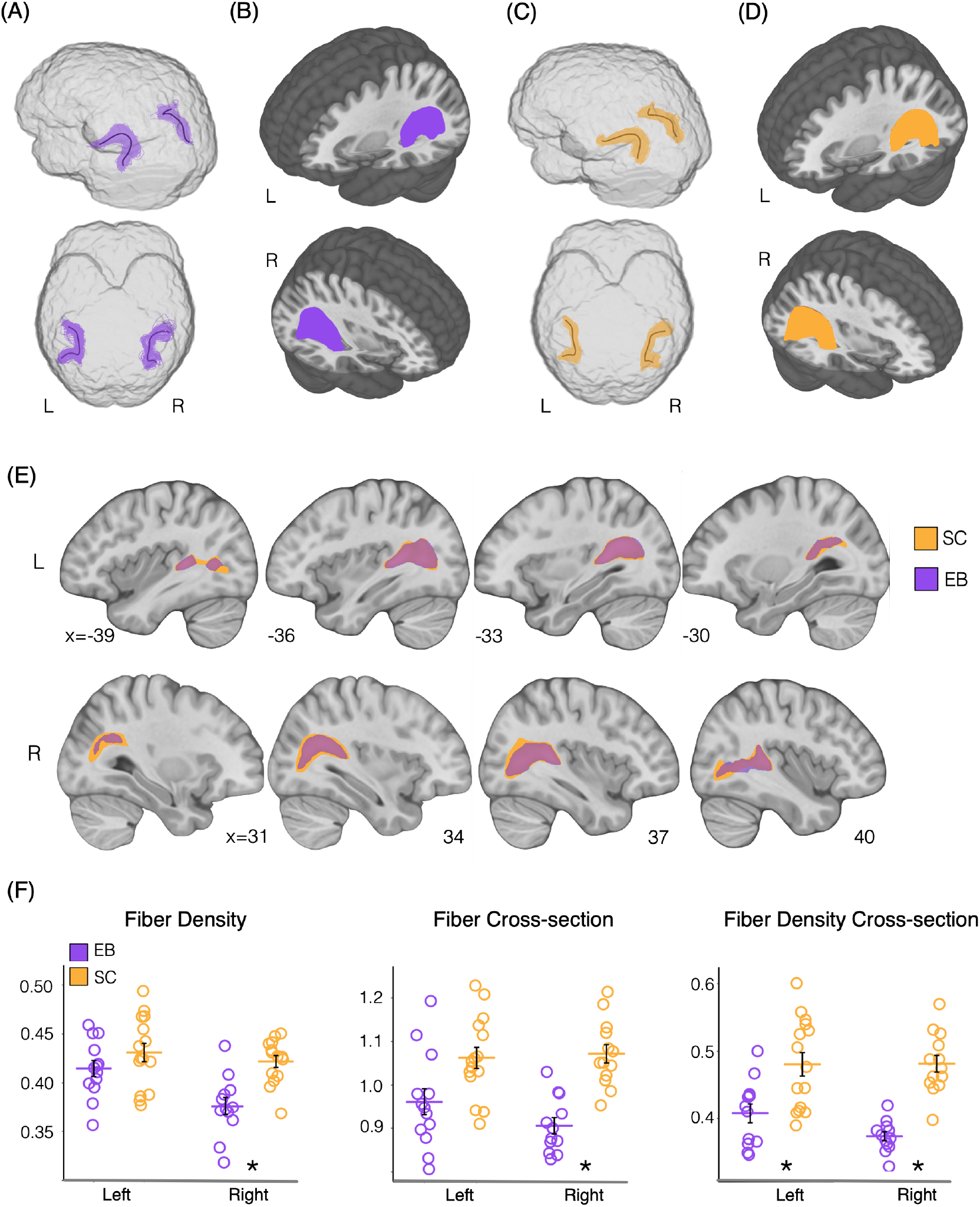
hMT+/V5 – hPT tractography and Whole Tract Analysis. **(A).** hMT+/V5 – hPT connections in a representative EB participant. Black line represents mean tract geometry. Tracts are shown in the template diffusion space. **(B).** Group-averaged structural pathways between hMT+/V5 and hPT for the EB group. Individual hMT+/V5 – hPT connections were binarized, overlaid and are shown at a threshold of >6 subjects. Results are depicted on the T1 MNI-152 template. **(C).** hMT+/V5 – hPT connections in a representative SC participant. **(D).** Group-averaged structural pathways between hMT+/V5 and hPT for the SC group. **(E).** Overlap of hMT+/V5 – hPT connections in the SC (orange) and the EB (purple) groups. Please note that the opacity of the tract in the EB blind group has been reduced in order to see the overlap. Results are depicted on the T1 MNI-152 template. **(F).** Dot plots represent the results of the whole tract analysis for early blinds (EB, in purple) and sighted controls (SC, in orange) for hMT+/V5 – hPT connections in the left and right hemisphere. Asterisks show significant (p < 0.05) differences in diffusion scalar values after Bonferroni correction for multiple comparisons. L: left, R: right.

The trajectory of the hMT+/V5 – hPT connections in the EB group was highly similar to that of the SC group (see Fig 4C). This was addressed by the Dice Similarity Coefficient. For the left hemisphere, the spatial overlap of the sum of binarized individual tract-density images was 0.84. For right hMT+/V5 – hPT connections, the spatial overlap between group-averaged tracts was equally high being the DSC value 0.80.

#### Whole tract analysis

The whole tract analysis revealed significant differences between EB end SC subjects in hMT+/V5 – hPT connections. We did not find differences between groups in the connectivity index (CI) (L: CI[SC] = 0.4 ± 0.1, CI[EB] = 0.4 ± 0.1, Paired t-Test, t(26) = 0.01, p = 0.9, d=0.01; R: CI[SC] = 0.4 ± 0.1, CI[EB] = 0.3 ± 0.1, Paired t-Test, t(26) = 1.52, p = 0.1, d=0.57). However, we did find differences between groups in diffusion scalar values, specifically in the right hemisphere. For left hMT+/V5 – hPT connections, we did not find differences between SCs and EBs in Fiber Density (FD) (L: FD[SC] = 0.43 ± 0.04, FD[EB] = 0.41 ± 0.03, Paired t-Test, t(26) = 1.31, p = 0.2, d=0.49) (see Fig 4F). For right hMT+/V5 – hPT connections, SCs showed higher FD values (M ± SD: 0.42 ± 0.02) than EBs (0.38 ± 0.03) (Paired t-Test, t(24) = 4.31, p = 2 x 10 ^-4^, d=1.67). We neither found differences between groups in Fiber Cross-section (FC) for left hMT+/V5 – hPT connections (L: FC[SC] = 1.06 ± 0.09, FC[EB] = 0.96 ± 0.11, Paired t-Test, t(26) = 2.62, p = 0.01, d=0.99). Similar to the results obtained for FD, SCs (M ± SD: 1.07 ± 0.08) showed higher FC values than EBs in right hMT+/V5 – hPT connections (0.91 ± 0.06) (Paired t-Test, t(23) = 5.89, p = 5 x 10 ^-6^, d=2.36). For the combined diffusion scalar, Fiber Density Cross-section (FDC), SCs presented higher values than EBs for hMT+/V5 – hPT connections in both hemispheres (L: FDC[SC] = 0.48 ± 0.07, FDC[EB] = 0.41 ± 0.05, Paired t-Test, t(25) = 3.14, p = 4 x 10^-3^, d=1.24; R: FDC[SC] = 0.48 ± 0.04, FDC[EB] =0.37 ± 0.02, Paired t-Test, t(23) = 7.48, p = 1 x 10 ^-7^, d=3.03. To note, when the lateralization of the group differences was tested using a 2×2 ANOVA (Group x Hemisphere), no interaction was found to be significant.

#### Along tract analysis

The along-tract analysis revealed the location of the between group differences at the sub-tract level. It confirmed hMT+/V5 – hPT connections in the right hemisphere to be more affected than its left counterpart. Whereas no differences between groups were found on FD and FC scalars for left hMT+/V5 – hPT connections, local differences in FDC were observed in the anterior parts of the tract (see upper row in Fig 5). For the right hMT+/V5 – hPT connections, structural effects for all diffusion scalars appear to be localized in posterior and central segments parts of the tract (see lower row in Fig 5).

**Figure 5.**
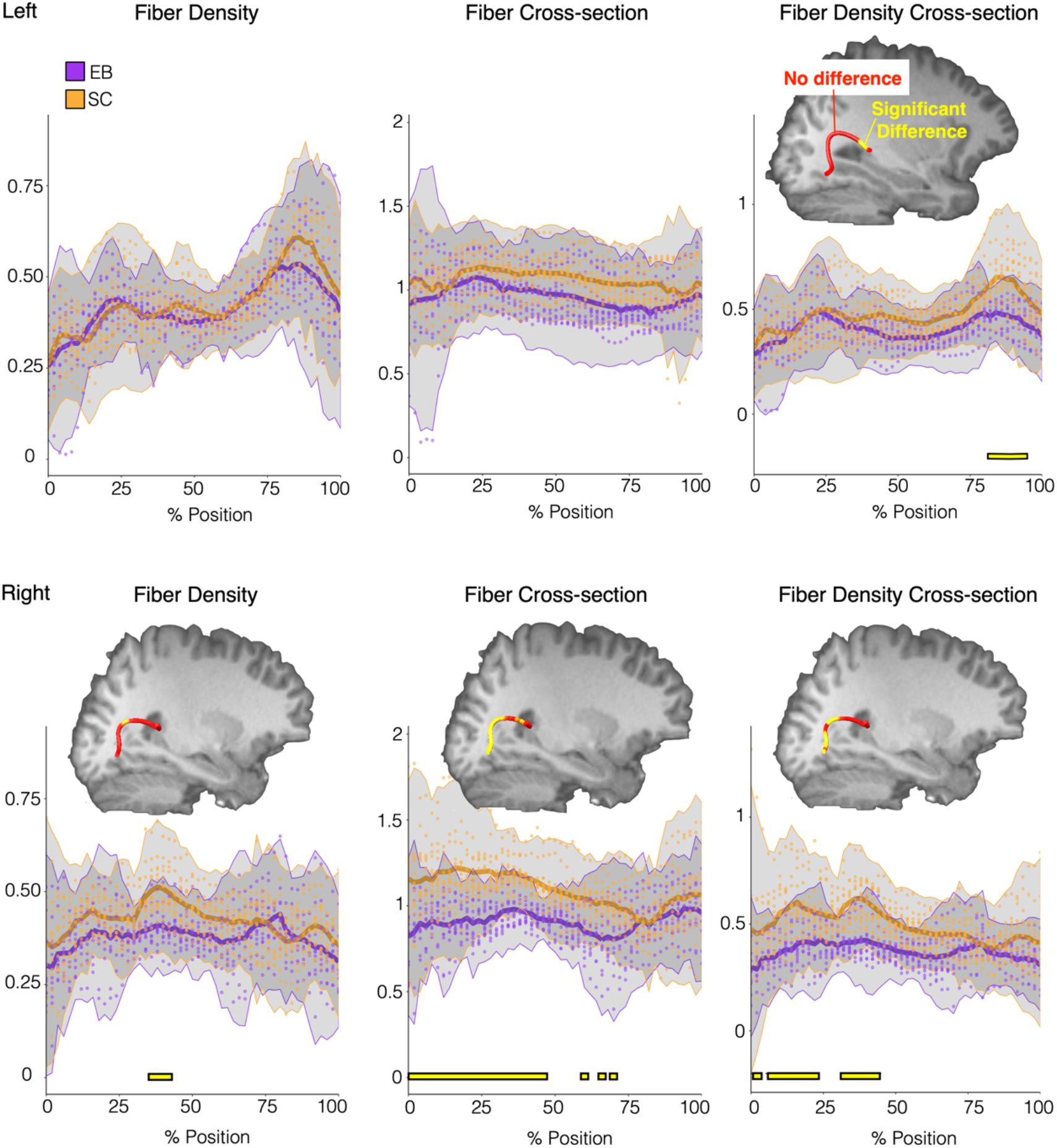
Along Tract Analysis results. Thick solid lines represent mean scalar values (Fiber Density, left; Fiber Cross-section, middle; Fiber Density Cross-section, right) for each group (EB in purple, SC in orange) for left (upper row) and right (lower row) hMT+/V5 – hPT connections. The shaded area illustrates values within ± 2.5 SD from the respective mean. Yellow lines in the lower part of each graph show the position along the tract where we find significant differences between groups (Bonferroni-corrected p < 0.05). Parts of the tract where significant differences between group are found, are also illustrated in the mean tract geometry of a representative subject. Yellow segments in the tract, represent sections where we found differences between groups, whereas red intervals represent segments without differences.

### Brain-Behavior Correlation

To explore whether the brain activity elicited by our moving and static sounds in hMT+/V5 and PT links to the ability of the listener to discriminate the direction and location of these sounds, we conducted between-subject correlations analyses. The multi-class decoding accuracies and beta parameter estimates were extracted from the peak coordinates of independently defined hMT+/V5 and PT regions. Behavioral accuracies recorded during fMRI data acquisition for discriminating motion directions (motion condition) and sound source locations (static condition) were correlated with decoding accuracies and beta parameter estimates separately. We also correlated microstructural diffusion values (FD, FC, and FDC) with behavioral performance. No significant correlation was observed between behavioral performance and the neural activity of hMT+/V5 and PT across groups. We did not find either any significant correlation between behavioral performance and microstructural diffusion values (full statistical report can be accessed at https://osf.io/7w8gu/).

## DISCUSSION

We adopted a multimodal imaging approach to investigate how early blindness alters the functional organization of the hMT+/V5 - hPT network for spatial hearing and the structural connectivity between those regions.

Whole-brain univariate analyses revealed that while both groups showed the strongest auditory motion selectivity in the bilateral superior temporal and precentral gyri, moving sounds also evoked preferential responses in a region anterior to hMT+/V5 in both sighted and blind individuals, a region previously termed hMTa (Rezk *et al*., 2020; Gurtubay-Antolin *et al*., 2021). hMTa overlaps with the anterior portion of hMT+/V5 (see Fig. 2C) suggesting that hMT+/V5 and hMTa are joined to form a continuous map. This observation relates to the evidence that category selective regions in vision are bound to a region just anterior to them showing amodal selectivity to the same category (Popham *et al*., 2021). Blind participants showed additional auditory motion selectivity in the posterior portion of hMT+/V5 suggesting that in the absence of sight, the posterior portion of hMT+/V5 -that is chiefly visual in sighted people-gets invaded by sounds coming from the anterior multisensory portion of hMT+/V5.

Going beyond univariate analyses, MVPA revealed motion direction and sound source location selective information in both hPT and hMT+/V5 of sighted and blind. The right hMT+/V5 showed enhanced decoding of auditory directions in the blind group. Interestingly, motion direction could also be decoded in the hMT+/V5 of sighted individuals, even if to a lower extent. These results contrast with previous studies finding no significant decoding of motion directions in hMT+/V5 of sighted people (Alink *et al*., 2012; Jiang, Stecker and Fine, 2014; Jiang *et al*., 2016). In those studies, however, directional selectivity was investigated exclusively in the horizontal axis, while the present study contained both horizontal and vertical auditory stimuli. This difference is crucial since we observed that activity patterns elicited in hMT+/V5 and hPT across the vertical and horizontal axes of motion differ to a much larger extent from activity patterns elicited by sounds within each axis of motion (Battal *et al*., 2019; Rezk *et al*., 2020). The dissimilarity analysis further supported an axis-of-motion organization and a separation between static and moving sounds in hPT and hMT+/V5 in both groups. In hPT, these results confirm previous observations for axis-of-motion (Battal et al., 2019). Interestingly, this organisation was weaker in the blind group compared to sighted. In hMT+/V5, the observation of an axis-of-motion organization for sounds are reminiscent of the columnar organization observed in vision (Albright, Desimone and Gross, 1984; Diogo *et al*., 2003), and the high-field fMRI studies suggesting a large-scale axis-of-motion organization in hMT+/V5 in humans (Zimmermann *et al*., 2011). This brings the resemblance between the coding of hMT+/V5 in vision and audition to an additional and finer-grained level (Kamitani and Tong, 2006), suggesting that the topographic organization principle of hMT+/V5 might be applied to the representation of auditory motion directions in blind and sighted people. The axis of auditory motion preference in hMT+/V5 in blind and sighted people suggests that the functional organization of hMT+/V5 for auditory directions is independent of visual experience.

In sighted individuals, even if hMT+/V5 shows preferential responses to visual motion, it also contains location selective representations of visual stimuli (Amano, Wandell and Dumoulin, 2009). If auditory information is being processed in hMT+/V5 of blind people using a computationally analog structure as the one observed in vision in sighted people, one may expect to find traces of sound source location in this region. In our study, we observed sound source location information in bilateral hMT+/V5 in EB, but not in the SC group. Our results therefore confirm and extend previous studies demonstrating that the right dorsal extrastriate occipital cortex in blind people contributes to spatial processing of sounds (Collignon et al. 2007; Collignon et al. 2009; Collignon et al. 2011; Collignon et al. 2009b).

Interestingly, we observed that the enhanced auditory tuning in hMT+/V5 in the blind co-occurs with a reduced decoding in hPT regions. The decreased decoding accuracies in the hPT for spatial hearing in the blind suggests that the absence of visual experience since birth not only influences the response properties of “visual” areas but also alters the functioning of the regions supporting the remaining senses. This re-distribution is not limited to auditory motion direction but is also observed for sound source location. It therefore seems that early visual deprivation triggers a network-level reorganization between occipital and temporal regions typically dedicated to spatial hearing.

Large-scale connectivity between separate sensory regions that are involved in related function could be a determining factor for the expression of crossmodal plasticity (Dormal and Collignon, 2011; Hannagan *et al*., 2015). Enhanced non-visual responses for moving stimuli observed in early blinds may therefore build on intrinsic connections between auditory and visual motion processing areas, which is supported by studies showing strong multisensory interactions between visual and auditory motion processing (Kitagawa and Ichihara, 2002). We recently demonstrated that hMT+/V5 contains shared neural representations for auditory and visual motion directions in sighted people (Rezk et al., 2020). We also found evidence for the existence of direct structural connection between hMT+/V5 and hPT in humans (Gurtubay-Antolin *et al*., 2021). Intrinsic anatomo-functional connections between visual and auditory motion-preferential regions could therefore play a crucial role in constraining the expression of crossmodal plasticity and re-distributing spatial hearing computation between hPT and hMT+/V5 in blindness. To further test this idea, we investigated the impact of visual deprivation on the structural connection between hPT and hMT+/V5 (Gurtubay-Antolin *et al*., 2021). Diffusion-weighted data in the EB and SC groups supported the potential existence of direct hMT+/V5 – hPT connections, with the trajectory and connectivity index of these projections being highly similar between SCs and EBs. These results speak in favor of the preservation of this pathway in visually deprived individuals. Our findings are in agreement with previous research conducted in opossums (Karlen, Kahn and Krubitzer, 2006) and humans (Shimony *et al*., 2005; Novello *et al*., 2018) that points to the overall preservation of structural connections in visual areas even in the absence of visual input since birth. We hypothesize that the preservation of the hMT+/V5 – hPT connections could play an important role in the multisensory integration of auditory and visual moving signals in sighted and in the expression of functionally specific crossmodal plasticity in the early blind group by constraining auditory moving sounds to specifically engage hMT+/V5 (Gurtubay-Antolin *et al*., 2021). It is however also possible that auditory information could reach hMT+/V5 through indirect pathways, for instance involving the parietal cortex (Bremmer et al., 2001, Rohe and Noppeney, 2016). Whereas we did not find macrostructural differences between groups in hMT+/V5 – hPT connections, whole-tract global white matter analyses showed decreased microstructural diffusion values (FD, FC and FDC) in EBs. Although the interpretation of diffusion-derived metrics remains controversial due to the indirect nature of the diffusion MRI measurement (Jones and Cercignani, 2010), FD and FC reductions have been related to axonal loss and atrophy of fiber morphology (reflective of the accumulated axon loss) (Raffelt et al., 2017). These results fit well with previous studies reporting reduced anisotropy values in visual pathways of EB individuals (Shimony *et al*., 2005; Ptito *et al*., 2008; Shu *et al*., 2009; Wang *et al*., 2013), which have been associated with deafferentation, demyelination, and neuronal degeneration (Beaulieu *et al*., 1996; Beaulieu, 2002). Alternatively, early visual deprivation could also lead to atypical myelination (Shimony et al., 2006). In this line, there is growing evidence that experience-dependent activity can induce myelination changes in the developing brain (Sampaio-Baptista and Johansen-Berg, 2017). The lack of visual input might have selectively affected the feedback projections connecting the hPT to hMT+/V5, which may play a role in audio-visual integration in sighted people (Ghazanfar and Schroeder, 2006). Since feedback projections appear late in the cortical development and have longer developmental times, they are more likely to be affected by sensory experience (Kral et al., 2017). The fact that this track losses its multisensory function to become mostly unimodal (auditory) in blind people may alter its microstructure (reducing FD and FC) in ways that are not yet well understood and would likely require studies with blind animals. To reveal which segments of the hMT+/V5 – hPT connections drive the differences between groups we implemented an along-tract analysis. Differences between groups emerged mostly in posterior and central parts of the right hMT+/V5 – hPT projections. No statistical difference between the reorganization across hemispheres emerged when formally tested. The higher alteration in posterior regions of the tracts might be explained by the higher dependence on visual input to fully develop.

In conclusion, our findings suggest that blindness triggers a network-level reorganization that enhances the recruitment of (posterior) hMT+/V5 in conjunction with a release in the computational workload of temporal regions (hPT) typically dedicated to spatial hearing. While visual experience does not affect the axis-of-motion organization in hMT+/V5, this functional organization in hPT region seemed to be weakened in EB. The structural connections between hMT+/V5 – hPT show similar macrostructure in both groups, despite microstructural alterations in these connections in visually deprived people.

## Acknowledgements

The project was funded in parts by an ERC starting grant MADVIS (Project: 337573) awarded to OC, the Belgian Excellence of Science (EOS) program (Project No. 30991544) awarded to OC, a Flagship ERA-NET grant SoundSight (FRS-FNRS PINT-MULTI R.8008.19) awarded to OC, and by the European Union’s Horizon 2020 research and innovation program under the Marie Skłodowska-Curie Grant Agreement No. 701250 awarded to VO. We would also like to express our gratitude to Marco Barilari, Stefania Benetti, Stephanie Cattoir who have helped with the data acquisition and Pietro Chiesa for continuing support with the auditory hardware. Computational resources have been provided by the supercomputing facilities of the Université catholique de Louvain (CISM/UCL) and the Consortium des Équipements de Calcul Intensif en Fédération Wallonie Bruxelles (CÉCI) funded by the Fond de la Recherche Scientifique de Belgique (F.R.S.-FNRS) under convention 2.5020.11 and by the Walloon Region. AG is supported by the Wallonie Bruxelles International Excellence Fellowship and the FSR Incoming PostDoc Fellowship by Université Catholique de Louvain. OC is a research associate; CB is postdoctoral researcher and MR is a research fellow at the Fond National de la Recherche Scientifique de Belgique (FRS-FNRS).

## REFERENCES

Albright, T. D., Desimone, R. and Gross, C. G. (1984) ‘Columnar organization of directionally selective cells in visual area MT of the macaque.’, Journal of neurophysiology, 51(1), pp. 16–31. doi: 6693933.

Alink, A. et al. (2012) ‘Auditory motion direction encoding in auditory cortex and high-level visual cortex.’, Human brain mapping, 33(4), pp. 969–78. doi: 10.1002/hbm.21263.

Allaire, J. J. (2015) ‘RStudio: Integrated development environment for R’, The Journal of Wildlife Management. doi: 10.1002/jwmg.232.

Amano, K., Wandell, B. A. and Dumoulin, S. O. (2009) ‘Visual Field Maps, Population Receptive Field Sizes, and Visual Field Coverage in the Human MT+ Complex’, Journal of Neurophysiology, 102(5), pp. 2704–2718. doi: 10.1152/jn.00102.2009.

Andersson, J. L. R. and Sotiropoulos, S. N. (2016) ‘An integrated approach to correction for off-resonance effects and subject movement in diffusion MR imaging’, NeuroImage, 125, pp. 1063–1078. doi: 10.1016/j.neuroimage.2015.10.019.

Avants, B. B. et al. (2008) ‘Symmetric diffeomorphic image registration with cross-correlation: Evaluating automated labeling of elderly and neurodegenerative brain’, Medical Image Analysis. doi: 10.1016/j.media.2007.06.004.

Battal, C. et al. (2019) ‘Representation of Auditory Motion Directions and Sound Source Locations in the Human Planum Temporale’, The Journal of Neuroscience, 39(12), pp. 2208–2220. doi: 10.1523/JNEUROSCI.2289-18.2018.

Battal, C. et al. (2020) ‘General Enhancement of Spatial Hearing in Congenitally Blind People’, Psychological Science, 31(9), pp. 1129–1139. doi: 10.1177/0956797620935584.

Baumgart, F. et al. (1999) ‘A movement-sensitive area in auditory cortex’, Nature, 400(6746), pp. 724–726. doi: 10.1038/23390.

Beaulieu, C. et al. (1996) ‘Changes in water diffusion due to Wallerian degeneration in peripheral nerve’, Magnetic Resonance in Medicine, 36(4), pp. 627–631. doi: 10.1002/mrm.1910360419.

Beaulieu, C. (2002) ‘The basis of anisotropic water diffusion in the nervous system – A technical review’, NMR in Biomedicine, 15(7–8), pp. 435–455. doi: 10.1002/nbm.782.

Behrens, T. E. J. et al. (2007) ‘Probabilistic diffusion tractography with multiple fibre orientations: What can we gain?’, NeuroImage, 34(1), pp. 144–155. doi: 10.1016/j.neuroimage.2006.09.018.

Benetti, S. et al. (2018) ‘White matter connectivity between occipital and temporal regions involved in face and voice processing in hearing and early deaf individuals’, NeuroImage, 179(March), pp. 263–274. doi: 10.1016/j.neuroimage.2018.06.044.

Benjamini, Y. and Yekutieli, D. (2001) ‘The control of the false discovery rate in multiple testing under dependency’, Annals of Statistics, 29(4), pp. 1165–1188. doi: 10.1214/aos/1013699998.

Bex, P. J., Simmers, A. J. and Dakin, S. C. (2003) ‘Grouping local directional signals into moving contours’, Vision Research, 43(20), pp. 2141–2153. doi: 10.1016/S0042-6989(03)00329-8.

Blank, H., Anwander, A. and von Kriegstein, K. (2011) ‘Direct Structural Connections between Voice-and Face-Recognition Areas’, Journal of Neuroscience, 31(36), pp. 12906–12915. doi: 10.1523/JNEUROSCI.2091-11.2011.

Blauert, J. (1997) Spatial Hearing : The Psychophysics of Human Sound Source Localization, 2nd ed, MIT Press. doi: 10.1109/jstqe.2002.804240.

Colby, J. B. et al. (2012) ‘Along-tract statistics allow for enhanced tractography analysis’, NeuroImage. doi: 10.1016/j.neuroimage.2011.11.004.

Collignon, O. et al. (2007) ‘Functional cerebral reorganization for auditory spatial processing and auditory substitution of vision in early blind subjects.’, Cerebral Cortex, 17(2), pp. 457–65. doi: 10.1093/cercor/bhj162.

Collignon, O. et al. (2009) ‘Reorganisation of the Right Occipito-Parietal Stream for Auditory Spatial Processing in Early Blind Humans. A Transcranial Magnetic Stimulation Study’, Brain Topography, 21(3–4), pp. 232–240. doi: 10.1007/s10548-009-0075-8.

Collignon, O. et al. (2011) ‘Functional specialization for auditory-spatial processing in the occipital cortex of congenitally blind humans’, Proceedings of the National Academy of Sciences, 108(11), pp. 4435–4440. doi: 10.1073/pnas.1013928108.

Cox, D. D. and Savoy, R. L. (2003) ‘Functional magnetic resonance imaging (fMRI) “brain reading”: Detecting and classifying distributed patterns of fMRI activity in human visual cortex’, NeuroImage, 19(2), pp. 261–270. doi: 10.1016/S1053-8119(03)00049-1.

Diogo, A. C. M. et al. (2003) ‘Electrophysiological imaging of functional architecture in the cortical middle temporal visual area of Cebus apella monkey.’, Journal of Neuroscience, 23(9), pp. 3881–3898. Available at: http://www.ncbi.nlm.nih.gov/pubmed/12736358.

Dormal, G. et al. (2016) ‘Auditory motion in the sighted and blind: Early visual deprivation triggers a large-scale imbalance between auditory and “visual” brain regions’, NeuroImage, 134(134), pp. 630–644. doi: 10.1016/j.neuroimage.2016.04.027.

Dormal, G. and Collignon, O. (2011) ‘Functional selectivity in sensory-deprived cortices.’, Journal of neurophysiology, 105(6), pp. 2627–30. doi: 10.1152/jn.00109.2011.

Eickhoff, S. B. et al. (2007) ‘Assignment of functional activations to probabilistic cytoarchitectonic areas revisited’, NeuroImage, 36(3), pp. 511–521. doi: 10.1016/j.neuroimage.2007.03.060.

Elbert, T. et al. (2002) ‘Expansion of the tonotopic area in the auditory cortex of the blind.’, The Journal of neuroscience : the official journal of the Society for Neuroscience, 22(22), pp. 9941–9944. doi: 22/22/9941 [pii].

Gougoux, F. et al. (2009) ‘Voice perception in blind persons: a functional magnetic resonance imaging study.’, Neuropsychologia, 47(13), pp. 2967–74. doi: 10.1016/j.neuropsychologia.2009.06.027.

Gurtubay-Antolin, A. et al. (2021) ‘Direct Structural Connections between Auditory and Visual Motion-Selective Regions in Humans’, The Journal of Neuroscience. doi: 10.1101/2020.06.11.145490.

Hannagan, T. et al. (2015) ‘Origins of the specialization for letters and numbers in ventral occipitotemporal cortex’, Trends in Cognitive Sciences, pp. 1–9. doi: 10.1016/j.tics.2015.05.006.

Hofman, P. M., Van Riswick, J. G. and Van Opstal, A. J. (1998) ‘Relearning sound localization with new ears.’, Nature neuroscience, 1(5), pp. 417–21. doi: 10.1038/1633.

Huk, A. C., Dougherty, R. F. and Heeger, D. J. (2002) ‘Retinotopy and functional subdivision of human areas MT and MST.’, The Journal of neuroscience : the official journal of the Society for Neuroscience, 22(16), pp. 7195–205. doi: 20026661.

Jeurissen, B. et al. (2014) ‘Multi-tissue constrained spherical deconvolution for improved analysis of multi-shell diffusion MRI data’, NeuroImage, 103, pp. 411–426. doi: 10.1016/j.neuroimage.2014.07.061.

Jiang, F. et al. (2016) ‘Early Blindness Results in Developmental Plasticity for Auditory Motion Processing within Auditory and Occipital Cortex’, Frontiers in Human Neuroscience, 10(July), p. 324. doi: 10.3389/fnhum.2016.00324.

Jiang, F., Stecker, G. C. and Fine, I. (2014) ‘Auditory motion processing after early blindness’, Journal of Vision, 14(13), pp. 4–4. doi: 10.1167/14.13.4.

Jones, D. K. and Cercignani, M. (2010) ‘Twenty-five pitfalls in the analysis of diffusion MRI data’, NMR in Biomedicine. doi: 10.1002/nbm.1543.

Kamitani, Y. and Tong, F. (2006) ‘Decoding seen and attended motion directions from activity in the human visual cortex.’, Current biology : CB, 16(11), pp. 1096–102. doi: 10.1016/j.cub.2006.04.003.

Karlen, S. J., Kahn, D. M. and Krubitzer, L. (2006) ‘Early blindness results in abnormal corticocortical and thalamocortical connections.’, Neuroscience, 142(3), pp. 843–58. doi: 10.1016/j.neuroscience.2006.06.055.

Kitagawa, N. and Ichihara, S. (2002) ‘Hearing visual motion in depth’, Nature, 416(6877), pp. 172–174. doi: 10.1038/416172a.

Krumbholz, K. et al. (2005) ‘Representation of interaural temporal information from left and right auditory space in the human planum temporale and inferior parietal lobe’, Cerebral Cortex, 15(3), pp. 317–324. doi: 10.1093/cercor/bhh133.

Majka, P. et al. (2019) ‘Unidirectional monosynaptic connections from auditory areas to the primary visual cortex in the marmoset monkey’, Brain Structure and Function, 224(1), pp. 111–131. doi: 10.1007/s00429-018-1764-4.

De Martino, F. et al. (2008) ‘Combining multivariate voxel selection and support vector machines for mapping and classification of fMRI spatial patterns’, NeuroImage, 43(1), pp. 44–58. doi: 10.1016/j.neuroimage.2008.06.037.

McFadyen, J., Mattingley, J. B. and Garrido, M. I. (2019) ‘An afferent white matter pathway from the pulvinar to the amygdala facilitates fear recognition’, eLife, 8, pp. 1–51. doi: 10.7554/elife.40766.

Middlebrooks, J. C. and Green, D. M. (1991) ‘Sound Localization by Human Listeners’, Annual Review of Psychology, 42(1), pp. 135–159. doi: 10.1146/annurev.ps.42.020191.001031.

Middlebrooks, J. C. and Green, D. M. (1992) ‘Observations on a principal components analysis of head-related transfer functions’, The Journal of the Acoustical Society of America, 92(1), pp. 597–599. doi: 10.1121/1.404272.

Møller, H. et al. (1996) ‘Binaural technique: Do we need individual recordings?’, AES: Journal of the Audio Engineering Society, 44(6), pp. 451–464.

Morris, D. M., Embleton, K. V. and Parker, G. J. M. (2008) ‘Probabilistic fibre tracking: Differentiation of connections from chance events’, NeuroImage, 42(4), pp. 1329– 1339. doi: 10.1016/j.neuroimage.2008.06.012.

Müller-Axt, C., Anwander, A. and von Kriegstein, K. (2017) ‘Altered Structural Connectivity of the Left Visual Thalamus in Developmental Dyslexia’, Current Biology, 27(23), pp. 3692–3698.e4. doi: 10.1016/j.cub.2017.10.034.

Musicant, A. D. and Butler, R. A. (1984) ‘The psychophysical basis of monaural localization’, Hearing Research, 14(2), pp. 185–190. doi: 10.1016/0378-5955(84)90017-0.

Nichols, T. et al. (2005) ‘Valid conjunction inference with the minimum statistic’, NeuroImage, 25(3), pp. 653–660. doi: 10.1016/j.neuroimage.2004.12.005.

Oosterhof, N. N., Connolly, A. C. and Haxby, J. V. (2016) ‘CoSMoMVPA: Multi-Modal Multivariate Pattern Analysis of Neuroimaging Data in Matlab/GNU Octave’, Frontiers in Neuroinformatics, 10(July), pp. 1–27. doi: 10.3389/fninf.2016.00027.

Palmer, S. M. and Rosa, M. G. P. (2006) ‘Quantitative analysis of the corticocortical projections to the middle temporal area in the marmoset monkey: Evolutionary and functional implications’, Cerebral Cortex, 16(9), pp. 1361–1375. doi: 10.1093/cercor/bhj078.

Park, H.-J. et al. (2007) ‘Reorganization of neural circuits in the blind on diffusion direction analysis.’, Neuroreport, 18(17), pp. 1757–60. doi: 10.1097/WNR.0b013e3282f13e66.

Pavani, F. et al. (2002) ‘A common cortical substrate activated by horizontal and vertical sound movement in the human brain’, Current Biology, 12(18), pp. 1584– 1590. doi: 10.1016/S0960-9822(02)01143-0.

Poirier, C. et al. (2005) ‘Specific activation of the V5 brain area by auditory motion processing: an fMRI study.’, Brain research. Cognitive brain research, 25(3), pp. 650– 8. doi: 10.1016/j.cogbrainres.2005.08.015.

Poirier, C. et al. (2006) ‘Auditory motion perception activates visual motion areas in early blind subjects’, NeuroImage, 31(1), pp. 279–85. doi: 10.1016/j.neuroimage.2005.11.036.

Popham, S. F. et al. (2021) ‘Visual and linguistic semantic representations are aligned at the border of human visual cortex’, Nature Neuroscience, 24(11), pp. 1628–1636. doi: 10.1038/s41593-021-00921-6.

Ptito, M. et al. (2008) ‘Alterations of the visual pathways in congenital blindness.’, Experimental brain research. Experimentelle Hirnforschung. Expérimentation cérébrale, 187(1), pp. 41–9. doi: 10.1007/s00221-008-1273-4.

Raffelt, D. et al. (2011) ‘Symmetric diffeomorphic registration of fibre orientation distributions.’, NeuroImage, 56(3), pp. 1171–80. doi: 10.1016/j.neuroimage.2011.02.014.

Raffelt, D., Tournier, J. D., Rose, S., et al. (2012) ‘Apparent Fibre Density: A novel measure for the analysis of diffusion-weighted magnetic resonance images’, NeuroImage, 59(4), pp. 3976–3994. doi: 10.1016/j.neuroimage.2011.10.045.

Raffelt, D., Tournier, J. D., Crozier, S., et al. (2012) ‘Reorientation of fiber orientation distributions using apodized point spread functions’, Magnetic Resonance in Medicine, 67(3), pp. 844–855. doi: 10.1002/mrm.23058.

Raffelt, D. et al. (2017) ‘Investigating white matter fibre density and morphology using fixel-based analysis’, NeuroImage, 144, pp. 58–73. doi: 10.1016/j.neuroimage.2016.09.029.

Rezk, M. et al. (2020) ‘Shared representation of visual and auditory motion directions in the human middle-temporal cortex’, Current Biology.

Rickham (1964) ‘World Medical Association. Code of Ethics of the World Medical Association. Declaration of Helsinki.’, British medical journal, 2, p. 177.

Saenz, M. et al. (2008) ‘Visual Motion Area MT+/V5 Responds to Auditory Motion in Human Sight-Recovery Subjects.’, The Journal of neuroscience : the official journal of the Society for Neuroscience, 28(20), pp. 5141–8. doi: 10.1523/JNEUROSCI.0803-08.2008.

Shimony, J. S. et al. (2005) ‘Diffusion Tensor Imaging Reveals White Matter Reorganization in Early Blind Humans’, Cerebral Cortex, 16(11), pp. 1653–1661. doi: 10.1093/cercor/bhj102.

Shu, N., et al. (2009) ‘Abnormal diffusion of cerebral white matter in early blindness.’, Human brain mapping, 30(1), pp. 220–7. doi: 10.1002/hbm.20507.

Smith, R. E. et al. (2012) ‘Anatomically-constrained tractography: Improved diffusion MRI streamlines tractography through effective use of anatomical information’, NeuroImage, 62(3), pp. 1924–1938. doi: 10.1016/j.neuroimage.2012.06.005.

Stelzer, J., Chen, Y. and Turner, R. (2013) ‘Statistical inference and multiple testing correction in classification-based multi-voxel pattern analysis (MVPA): Random permutations and cluster size control’, NeuroImage, 65, pp. 69–82. doi: 10.1016/j.neuroimage.2012.09.063.

Tootell, R. B. et al. (1995) ‘Functional analysis of human MT and related visual cortical areas using magnetic resonance imaging’, J.Neurosci., 15(0270-6474 (Print)), pp. 3215–3230.

Tournier, J. D., Calamante, F. and Connelly, A. (2007) ‘Robust determination of the fibre orientation distribution in diffusion MRI: Non-negativity constrained super-resolved spherical deconvolution’, NeuroImage, 35(4), pp. 1459–1472. doi: 10.1016/j.neuroimage.2007.02.016.

Tournier, J. D., Calamante, F. and Connelly, A. (2010) ‘Improved probabilistic streamlines tractography by 2nd order integration over fibre orientation distributions’, Proceedings of the International Society for Magnetic Resonance in Medicine, 88(2003), p. 2010.

Tournier, J. D., Calamante, F. and Connelly, A. (2012) ‘MRtrix: Diffusion tractography in crossing fiber regions’, International Journal of Imaging Systems and Technology, 22(1), pp. 53–66. doi: 10.1002/ima.22005.

Tschentscher, N. et al. (2019) ‘Reduced structural connectivity between left auditory thalamus and the motion-sensitive planum temporale in developmental dyslexia’, Journal of Neuroscience, 39(9), pp. 1720–1732. doi: 10.1523/JNEUROSCI.1435-18.2018.

Tustison, N. J. et al. (2010) ‘N4ITK: Improved N3 bias correction’, IEEE Transactions on Medical Imaging, 29(6), pp. 1310–1320. doi: 10.1109/TMI.2010.2046908.

Veraart, J. et al. (2016) ‘Denoising of diffusion MRI using random matrix theory.’, NeuroImage, 142, pp. 394–406. doi: 10.1016/j.neuroimage.2016.08.016.

Wang, D. et al. (2013) ‘Altered white matter integrity in the congenital and late blind people’, Neural Plasticity, 2013(1). doi: 10.1155/2013/128236.

Warren, J., Zielinski, B. and Green, G. (2002) ‘Perception of sound-source motion by the human brain’, Neuron, 34, pp. 139–148. Available at: http://www.sciencedirect.com/science/article/pii/S0896627302006372 (Accessed: 7 February 2014).

Watson, J. D. G. et al. (1993) ‘Area V5 of the Human Brain: Evidence from a Combined Study Using Positron Emission Tomography and Magnetic Resonance Imaging’, Cereb. Cortex, 3(2), pp. 79–94. doi: 10.1093/cercor/3.2.79.

Wightman, F. L. and Kistler, D. J. (1989) ‘Headphone simulation of free-field listening. II: Psychophysical validation’, The Journal of the Acoustical Society of America, 85(2), pp. 868–878. doi: 10.1121/1.397558.

Wolbers, T., Zahorik, P. and Giudice, N. a (2011) ‘Decoding the direction of auditory motion in blind humans.’, NeuroImage, 56(2), pp. 681–7. doi: 10.1016/j.neuroimage.2010.04.266.

Worsley, K. J. et al. (1996) ‘A unified statistical approach for determining significant signals in images of cerebral activation’, Human Brain Mapping, 4(1), pp. 58–73. doi: 10.1002/(SICI)1097-0193(1996)4:1<58::AID-HBM4>3.0.CO;2-O.

Zeki, S. et al. (1991) ‘A direct demonstration of functional specialization in human visual cortex’, J.Neurosci., 11(3), pp. 641–649.

Zeng, H. and Constable, R. T. (2002) ‘Image distortion correction in EPI: Comparison of field mapping with point spread function mapping’, Magnetic Resonance in Medicine, 48(1), pp. 137–146. doi: 10.1002/mrm.10200.

Zhang, Y., Brady, M. and Smith, S. (2001) ‘Segmentation of brain MR images through a hidden Markov random field model and the expectation-maximization algorithm’, IEEE Transactions on Medical Imaging. doi: 10.1109/42.906424.

Zimmermann, J. et al. (2011) ‘Mapping the Organization of Axis of Motion Selective Features in Human Area MT Using High-Field fMRI’, PLoS ONE. Edited by H. P. Op de Beeck, 6(12), p. e28716. doi: 10.1371/journal.pone.0028716.

